# Biomolecular condensates amplify mRNA decapping by coupling protein interactions with conformational changes in Dcp1/Dcp2

**DOI:** 10.1101/2020.07.09.195057

**Authors:** Ryan W. Tibble, Anaïs Depaix, Joanna Kowalska, Jacek Jemielity, John D. Gross

## Abstract

Cells organize biochemical processes into biological condensates. P-bodies are cytoplasmic condensates enriched in factors important for mRNA degradation. P-bodies have been identified as sites of both mRNA storage and decay, but how these opposing outcomes may be achieved in condensates is unresolved. A critical step in mRNA degradation is removal of the 5’-7-methylguanosine cap by Dcp1/Dcp2, which is highly enriched in P-bodies. Dcp1/Dcp2 activity is repressed in condensates *in vitro* and requires the activator Edc3. Activation of decapping is amplified in condensates relative to the surrounding solution due to stabilization of an autoinhibited state in Dcp1/Dcp2. Edc3 couples a conformational change in the Dcp1/Dcp2 active site with alteration of the protein-protein interactions driving phase separation to activate decapping in condensates. The composition-dependent regulation of enzyme activity in condensates occurs over length scales ranging from microns to Ångstroms and may control the functional state of P-bodies and related phase-separated compartments.

**HIGHLIGHTS:**

- mRNA decapping in droplets is repressed
- Catalytically inert droplets are activated by a change in condensate composition
- A switch in enzymatic activity requires a conformational change in condensates
- Condensates amplify enzyme activation compared to surrounding solution

## INTRODUCTION

Cellular mRNA is highly regulated from its initial transcription to eventual degradation and much of this regulation arises from coordinated interactions with proteins (Moore, 2005). The assembly and organization of messenger ribonucleoprotein (mRNP) complexes has functional consequences on the transcript, dictating whether it is spliced, exported from the nucleus, actively translated, or degraded. It has become increasingly appreciated that many mRNPs accumulate in biomolecular condensates under normal and stressed conditions. Splicing and pre-mRNA processing factors are localized to nuclear Cajal bodies and paraspeckles, for example, while proteins involved in translational repression and degradation are enriched in cytoplasmic stress granules (SGs) and P-bodies (PBs), respectively (Courchaine et al., 2016; Decker and Parker, 2012; Müller-McNicoll and Neugebauer, 2013).

Biomolecular condensates have emerged as a mechanism to control numerous cellular reactions in distinct ways: partitioning proteins to regulate productive interactions, increasing the local concentration to accelerate enzymatic activity, buffering of activity to prevent aberrant transcription, and thermodynamically coupling interactions to enforce directionality in ribosome assembly (Banani et al., 2016; Gallego et al., 2020; Hnisz et al., 2017; Riback et al., 2020; Sheu-Gruttadauria and MacRae, 2018). The emergent properties afforded by formation of an extensive interaction network in condensates also have the potential to enhance enzymatic activity beyond local concentration effects (Banani et al., 2017). Such enzymatic enhancements could arise from the coupling of enzyme allostery to interactions that span the diameter of condensates. How the collective properties across length scales are coupled to enzyme catalysis is poorly understood.

P-bodies are a conserved class of phase-separated biomolecular condensates present under normal cellular conditions. Components of bulk 5’-3’ mRNA decay represent the most well-conserved and abundant proteins in P-bodies and, together, these proteins serve to form a dense network of redundant interactions that allow for P-body formation (Franks and Lykke-Andersen, 2008; Hubstenberger et al., 2017; Parker and Sheth, 2007; Rao and Parker, 2017; Teixeira and Parker, 2007). Although much progress has been made in identifying the molecular composition and interactions within P-bodies, the biological function of PBs remains unresolved. In particular, it is unclear whether P-bodies promote active mRNA processing or are sites of storage. Several studies suggest P-bodies are sites of mRNA decay due to the accumulation of mRNA and decay intermediates in P-bodies when 5’-3’ degradation factors are knocked-out or mutated (Chan et al., 2018; Mugler et al., 2016; Sheth and Parker, 2003). Alternatively, multiple studies propose P-bodies function primarily as sites of mRNA storage due to the absence of decay intermediates in sequencing and live cell imaging data (Horvathova et al., 2017; Hubstenberger et al., 2017; Tutucci et al., 2018). Moreover, mRNAs exhibited context-dependent degradation in P-bodies and can be restored to the translating pool upon recovery from cell stress (Aizer et al., 2014; Brengues et al., 2005; Wang et al., 2018). Given their heterotypic and dynamic nature, however, isolating and studying P-bodies has been challenging and presents a current hurdle in elucidating their biological function.

The assembly of P-bodies, similar to other biomolecular condensates, relies on a network of multivalent interactions between resident proteins often mediated by intrinsically disordered regions (IDRs) (Boeynaems et al., 2018; Hyman et al., 2014). Many factors associated with 5’-3’ decay contain structured domains flanked by IDRs important for recruitment to P-bodies, interactions with RNA, and other decay protein cofactors (Jonas and Izaurralde, 2013). This includes the conserved decapping complex, comprised of the catalytic subunit Dcp2 and its obligate activator Dcp1, which is responsible for hydrolysis of the 7-methylguanosine (m7G) cap from mRNA (Arribas-Layton et al., 2013; Beelman et al., 1996; Dunckley and Parker, 1999; Wang et al., 2002). Decapping is generally considered the irreversible step in 5’-3’ RNA decay, committing the transcript to rapid degradation, and represents a critical checkpoint in regulating mRNA turnover (Mugridge et al., 2018; Nagarajan et al., 2013). In addition to being a major component of cellular P-bodies, Dcp2 function has been implicated in spinal muscular atrophy, innate immunity, and the interferon response (Abernathy and Glaunsinger, 2015; Li et al., 2012; Shukla and Parker, 2014). Mammalian Dcp1 is cleaved by poliovirus, leading to its degradation and disruption of P-bodies (Dougherty et al., 2011).

A disordered C-terminus in yeast Dcp2 and metazoan Dcp1 is critical for their localization to P-bodies. Moreover, excision of the C-terminus in yeast leads to dysregulation of numerous transcripts and conditional lethality (Fromm et al., 2014; He et al., 2018). The localization and regulation imparted by disordered regions in the decapping complex is mediated through conserved short-linear interaction motifs (Fromm et al., 2014; Jonas and Izaurralde, 2013; Xing et al., 2018). Recently, positive and negative regulatory motifs in the disordered C-terminus of Dcp2 were identified in yeast Dcp2 that affect catalytic activity *in vitro* and *in vivo* (He and Jacobson, 2015; Paquette et al., 2018). These results established a model where Dcp2 is autoinhibited and coactivators alleviate autoinhibition to increase RNA binding and catalysis (Lobel et al., 2019; Paquette et al., 2018). However, the molecular mechanisms for regulation of autoinhibition are not well understood.

Edc3 is an activator of decapping responsible for the degradation of specific mRNA transcripts and regulation of deadenylation-independent 5’-3’ decay in budding yeast (Badis et al., 2004; He et al., 2018, 2014). Edc3 is also required for proper neuronal development as mutations disrupting activity lead to developmental disabilities in humans (Ahmed et al., 2015). Edc3 interacts with positive regulatory motifs in the C-terminus of Dcp2 to alleviate autoinhibition and promote Dcp2 activity (Fromm et al., 2012; Harigaya et al., 2010; He and Jacobson, 2015; Paquette et al., 2018). Edc3 localizes to P-bodies and has been shown to form multivalent interactions with RNA and the C-terminus of Dcp2 to promote liquid-liquid phase separation (Damman et al., 2019; Fromm et al., 2014; Schütz et al., 2017). Initial studies suggested Edc3-mediated phase separation of Dcp1/Dcp2 inhibited the rate of decapping (Schütz et al., 2017). These studies, however, indirectly measured decapping activity in condensates and a direct investigation of decapping in condensates has not been carried out. As a result, it is unclear how phase separation regulates decapping.

In this study, we address the relationship between molecular organization and function by reconstituting minimal decapping condensates containing Dcp1/Dcp2 and the activator Edc3. Using a dual-labeled substrate for simultaneous monitoring of the 5’-cap and RNA body, we directly measure rates of decapping in condensates by fluorescence microscopy. We find Dcp1/Dcp2 sequestered in condensates is inactive. In contrast, addition of Edc3 reorganizes the underlying network of interactions in droplets and cooperatively enhances activity of Dcp1/Dcp2 by 80-fold. The fold activation of decapping in droplets by Edc3 is greater than in solution, not because enzyme turnover is accelerated, but instead due to increased repression of activity when Edc3 is absent. We show that a conformational change in Dcp1/Dcp2 is promoted by Edc3 to activate decapping and suggest that in the absence of Edc3, the droplet environment can further bias a conformational equilibrium in Dcp1/Dcp2 to allow for robust repression of enzyme activity. Our findings suggest droplet composition can tune conformational dynamics of enzymes to affect activity, which may be an emergent property of biomolecular condensates that is used for controlling RNA degradation and other biochemical reactions in cells.

## RESULTS

### Liquid-liquid phase separation of Dcp1/Dcp2 is potentiated by Edc3

Dcp2 contains well-structured regulatory (NRD) and catalytic (CD) core domains at its N-terminus that undergo conformational changes during the catalytic cycle (Dcp2_core_ in **Figure 1A**). Recently, we reconstituted a construct of *S. pombe* Dcp2 containing a portion of the C-terminal IDR necessary for regulation of decapping (Dcp2_ext_ in **Figure 1A**) (Paquette et al., 2018). In the absence of activators, Dcp1/Dcp2_ext_ is autoinhibited by inhibitory motifs (IMs) in the C-terminal IDR. Edc3 interacts with flanking helical leucine rich motifs (HLMs) in the IDR to alleviate autoinhibition (**Figure 1A** and **1B**). These experiments on Dcp1/Dcp2_ext_ were carried out with soluble protein, and we noticed at higher concentrations Dcp1/Dcp2_ext_ underwent phase separation (**Figure 1C**, top row). Because prior studies examined Dcp1/Dcp2 condensates using a construct lacking IMs, we characterized how these elements of the C-terminus contribute to phase separation of Dcp1/Dcp2_ext_ in parallel with those formed in complex with Edc3 (Fromm et al., 2014; Schütz et al., 2017).

**Figure 1.**
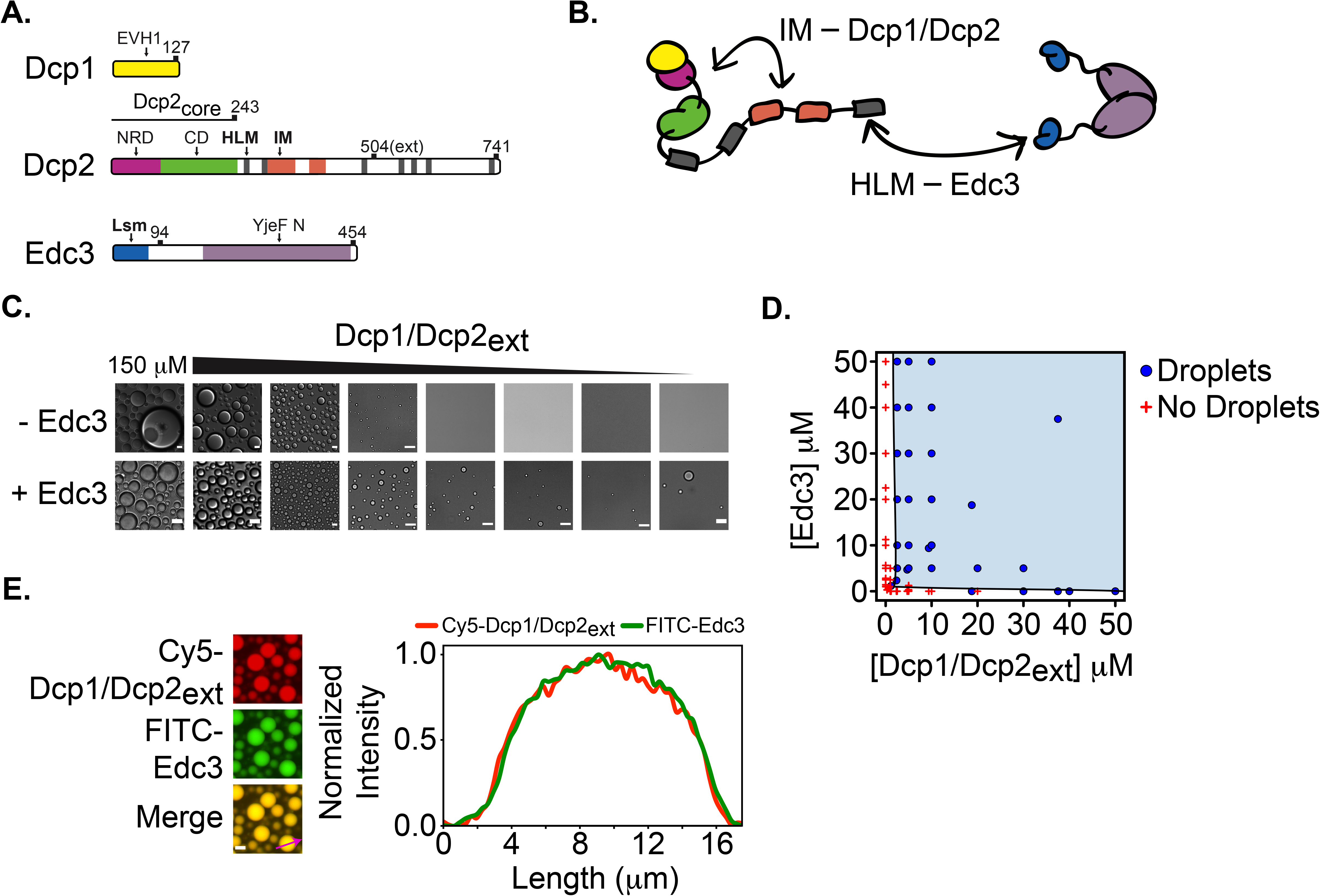
Edc3 enhances Dcp1/Dcp2_ext_ phase separation. **A**. Schematic for Dcp1, Dcp2, and Edc3. Domains and motifs are labeled and white space is indicative of disordered regions. Dcp2 contains structured core domains, Dcp2_core_, and the construct used in this study, Dcp2_ext_, contains regulatory elements in the C-terminal IDR. **B**. Cartoon of Dcp1/Dcp2_ext_ and Edc3 highlighting the proposed interaction between the IMs and core domains of Dcp1/Dcp2_ext_ and between HLMs and Edc3. Cartoons are colored as shown in A. **C**. Dcp1/Dcp2_ext_ undergoes liquid-liquid phase separation and addition of stoichiometric amounts of Edc3 reduces the critical concentration for phase separation twenty-fold. **C**. Phase diagram of Dcp1/Dcp2_ext_ and Edc3 *in vitro* phase separation (teal region). **D**. Dcp1/Dcp2_ext_ and Edc3 are equally enriched and homogenously distributed in droplets. Protein concentration is 50 μM. Scale bar, 10 μm.

Droplets containing Dcp1/Dcp2_ext_ varied in cross-sectional area from less than 50 μm^2^ to greater than 500 μm^2^ in diameter and underwent fusion, indicative of a dynamic, liquid-like biomolecular condensate (**Figure S1A**, and **S1E**). Dcp2_ext_ and residues 274-504 of the Dcp2 C-terminus also underwent concentration-dependent phase separation, indicating low-complexity regions in Dcp2 are sufficient to promote phase separation, which is in agreement with in vivo studies showing the Dcp2 C-terminus is required for P-body recruitment (**Figure S1C** and **S1D**) (Fromm et al., 2014). Thus, our data demonstrates Dcp1/Dcp2_ext_ can form large, microscopic condensates mediated by the C-terminus.

Because Edc3 can form multivalent interactions with the HLMs in the Dcp2 C-terminus, we tested how Edc3 influenced the critical concentration for phase separation. Edc3 reduced the critical concentration of Dcp1/Dcp2_ext_ condensates twenty-fold (**Figure 1C** bottom row, and **1D**). Fluorescently labeled Dcp1/Dcp2_ext_ and Edc3 colocalized to droplets and were homogenously distributed (**Figure 1E**). Similar to Dcp1/Dcp2_ext_ droplets, Dcp1/Dcp2_ext_/Edc3 droplets undergo fusion and are concentration-dependent; however, droplets containing Edc3 are not larger than 100 μm^2^ in cross-sectional area, which is in contrast to the numerous droplets with cross-sectional area greater than 100 μm^2^ observed for Dcp1/Dcp2_ext_ (**Figures 1B, S1C**, and **S1E**). The reduction in size distribution and concomitant lower critical concentration suggests Edc3 contributes to the network of protein interactions driving phase separation of Dcp1/Dcp2.

### The interactions required for Dcp1/Dcp2_ext_ and Dcp1/Dcp2_ext_/Edc3 condensation are different

The critical concentration of Dcp1/Dcp2_ext_ condensate formation depends on Edc3, raising the possibility the interactions promoting droplet formation differ. Short-linear interaction motifs typically mediate phase separation, and we hypothesized different motifs in the C-terminus of Dcp2 may contribute to condensation of Dcp1/Dcp2_ext_ in the presence or absence of Edc3 (**Figure 2A**). If phase separation is driven by interactions between IMs and the structured regions of Dcp1/Dcp2_ext_ *in trans*, then removal of IMs should abrogate phase separation. However, if Edc3 results in a molecular reorganization of droplets to favor interactions between Edc3 and HLMs in Dcp2, then phase separation of Dcp1/Dcp2/Edc3 should be independent of IMs (**Figure 2A**). To test these models, we developed a series of truncations in Dcp2 and Edc3 and looked for the presence of droplets using microscopy (**Figure 2B**).

**Figure 2.**
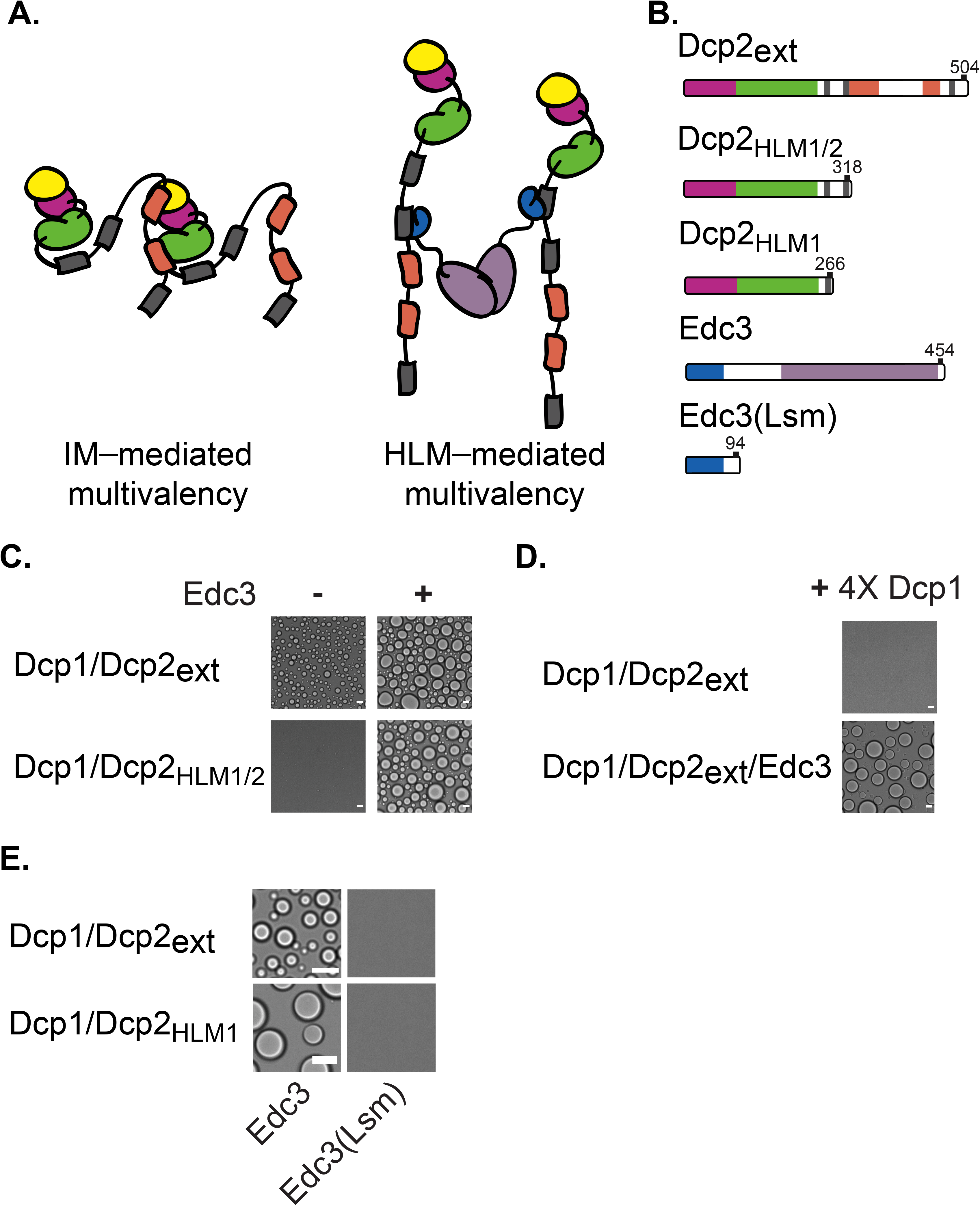
Interactions underlying Dcp1/Dcp2_ext_ and Dcp1/Dcp2_ext_/Edc3 phase separation are distinct. **A**. Interaction between IMs of one Dcp2 molecule and a neighboring Dcp1/Dcp2_ext_ *in trans* could mediate Dcp1/Dcp2_ext_ phase separation whereas Dcp1/Dcp2_ext_/Edc3 phase separation is promoted by Edc3 interaction with HLMs across multiple Dcp2 molecules. **B**. Dcp2 truncations used to determine interactions important for phase separation. **C**. Removal of inhibitory motifs from the C-terminus of Dcp2 ablates phase separation of the decapping complex yet retains the ability undergo droplet formation upon addition of Edc3. Concentration of Dcp1/Dcp2_HLM1/2_ is 300 μM, Dcp1/Dcp2_ext_ is 100 μM, Dcp1/Dcp2_HLM1/2_/Edc3 and Dcp1/Dcp2_ext_/Edc3 are at 50 μM **D**. Addition of four-fold molar excess of Dcp1 disrupts Dcp1/Dcp2_ext_ phase separation (top) but not Dcp1/Dcp2_ext_/Edc3 droplets (bottom). Concentration of Dcp1 is 200 μM, Dcp1/Dcp2 and Dcp1/Dcp2/Edc3 is 50 μM. **E**. Dimerization of Edc3 is necessary to promote phase separation of Dcp1/Dcp2 containing a single (Dcp1/Dcp2_HLM1_, top right) or multiple HLMs (bottom right). Dcp1/Dcp2_HLM1_ and Edc3 were stoichiometrically mixed at 150 μM. Dcp1/Dcp2_ext_ and Edc3 were mixed at 50 μM.

A construct lacking IMs, Dcp2_HLM1/2_, abolished phase separation of Dcp1/Dcp2 but not Dcp1/Dcp2/Edc3 (**Figure 2C**). We previously demonstrated an IM interacts with Dcp1 *in trans* and a region of the Dcp2 C-terminus can interact with Dcp1 *in cis* (Paquette et al., 2018; Wurm et al., 2016). Thus, Dcp1 could contribute to multivalency in Dcp1/Dcp2_ext_ phase separation. This predicts excess Dcp1 in *trans* should compete for Dcp1—Dcp2 intermolecular interactions and prevent phase separation of Dcp1/Dcp2_ext_ droplets, but not affect Edc3-driven phase separation. To test this possibility, we added four-fold excess Dcp1 to Dcp1/Dcp2_ext_ condensates and did not observe the formation of liquid droplets (**Figure 2D**). Addition of Edc3, however, still resulted in the formation of droplets in the presence of excess Dcp1 and Edc3-dependent phase separation required Edc3 dimerization (**Figure 2D** and **2E**). Thus, Dcp2 IMs and Dcp1 are important for phase separation of Dcp1/Dcp2_ext_ while Dcp2 HLMs and Edc3 are important for droplet formation of Dcp1/Dcp2_ext_/Edc3. These results suggest condensate composition can lead to changes in the network of underlying molecular interactions critical for their formation.

### Edc3 activates decapping in droplets

Since the molecular organization of Dcp1/Dcp2_ext_ and Dcp1/Dcp2_ext_/Edc3 droplets involves interactions important for autoinhibition and activation, respectively, we asked whether condensates could exhibit different decapping activity. To test and quantify decapping by Dcp1/Dcp2_ext_ in droplets, we synthesized a capped RNA substrate containing a 5’ cap conjugated to fluorescein and 3’ adenosine conjugated to Cy5 (**Figures 3A** and **S2**). This dual-labeled substrate allows for simultaneous monitoring of the 5’-cap and RNA body, making it possible to directly study decapping in condensates by fluorescence microscopy. We first introduced this probe to Dcp1/Dcp2_ext_ condensates and observed its enrichment in droplets and monitored fluorescence from the cap and RNA over time (**Figure 3B**). After twenty minutes we did not notice an appreciable loss in fluorescence intensity, suggesting Dcp1/Dcp2_ext_ in these droplets decaps RNA at a rate less than 0.006 min^−1^ (**Figure 3C**). However, upon addition of the dually labelled probe to droplets formed in a solution of Dcp1/Dcp2_ext_ and excess Edc3, a rapid, time-dependent decrease in fluorescein fluorescence was observed, indicating loss of cap from the droplets (**Figure 3D**). The rate of m^7^GDP loss in Dcp1/Dcp2_ext_/Edc3 droplets was 0.56 min^−1^ and we estimate Edc3 enhanced decapping in droplets 80-fold (**Figure 3E**). Localization of the RNA body, as monitored by Cy5 fluorescence, did not greatly change over time and suggests it is retained within condensates despite loss of the cap. These observations are dependent on enzyme catalysis as omission of magnesium ions results in an inappreciable change in signal for 5’ cap and RNA body within Dcp1/Dcp2_ext_/Edc3 droplets (**Figure S2E** and **S2F**). We conclude decapping can occur within droplets but critically depends on the presence of the activator Edc3.

**Figure 3.**
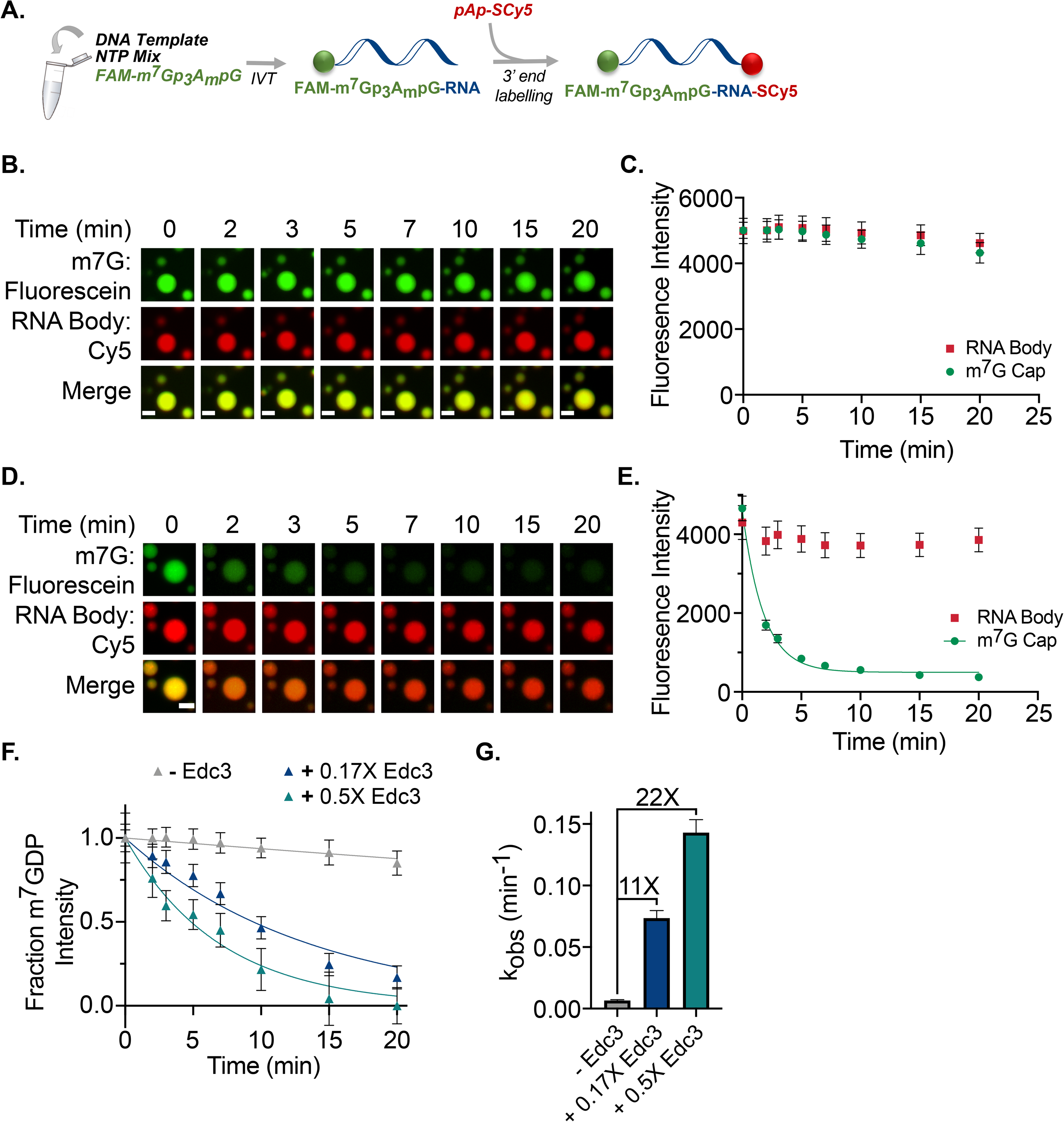
The ability of phase-separated Dcp1/Dcp2_ext_ to decap RNA is modulated by Edc3. **A**. Synthesis scheme for two-color fluorescent 35mer RNA capped with fluorescein m^7^G cap (FAM-m^7^GDP) and labeled at the 3’ end with Cy5-adenosine (see Methods). **B, C**. Addition of dual-labeled capped RNA to Dcp1/Dcp2_ext_ (60 μM total protein concentration) droplets shows minimal signal loss in both FAM-m^7^GDP and Cy5-RNA over twenty minutes. **D, E**. Addition of dual-labeled, capped RNA to 5 μM Dcp1/Dcp2_ext_ and 80 μM Edc3 droplets causes rapid time-dependent loss of m^7^GDP signal from the droplet while the RNA body is retained. **F, G**. Addition of substoichiometric concentrations of Edc3 to preformed Dcp1/Dcp2_ext_ droplets (60 μM total Dcp1/Dcp2_ext_ concentration) results in dose-dependent loss of FAM-m^7^GDP signal from droplets. Intensities in C, E, and G represent the mean value for ten droplets. Error bars represent s.e.m. Scale bar, 5 μm.

Since Dcp1/Dcp2_ext_ droplets are inhibited in their ability to decap mRNA substrate, we tested if the addition of Edc3 to these droplets is sufficient to activate decapping. Dcp1/Dcp2_ext_ droplets were pre-formed in the presence of dual-labeled RNA substrate and Edc3 was added just prior to initiation of the reaction with Mg^2+^. Substoichiometric concentrations of Edc3 resulted in a time-dependent loss of m^7^GDP signal from within the droplet at least 22-fold greater than droplets containing only Dcp1/Dcp2_ext_ (**Figure 3F** and **3G**). Together, these data indicate Edc3 is recruited to Dcp1/Dcp2_ext_ droplets to activate decapping and Dcp1/Dcp2_ext_ sequestered in the condensed phase is sensitive to alteration in droplet composition.

Our results show Edc3 can activate decapping 80-fold in droplets. This suggests Dcp1/Dcp2_ext_/Edc3 condensates may be sites of increased decapping activity relative to the surrounding solution. Alternatively, Dcp1/Dcp2_ext_ condensates may be repressed in activity relative to the surrounding solution. To distinguish these possibilities, we measured the maximal rate of decapping of the dual-labeled substrate in dilute phase using Dcp1/Dcp2_core_, which is not autoinhibited and does not undergo phase separation (Paquette et al., 2018). The maximal decapping rate observed in Dcp1/Dcp2_ext_/Edc3 droplets does not differ greatly from that observed for Dcp1/Dcp2_core_ (**Figure S3**). However, the rate of decapping determined in Dcp1/Dcp2_ext_ droplets is at least 30-fold slower than for Dcp1/Dcp2_core_. Thus, we conclude the droplet environment does not provide conditions favoring increased decapping activity by Edc3, but rather, a greater degree of repression in the absence of Edc3. Furthermore, our results suggest a high concentration of Dcp1/Dcp2_ext_ in droplets per se is not sufficient for active decapping but instead requires binding to the activator Edc3.

### Activation of decapping by Edc3 is coupled to phase separation

In order to differentiate further between enzyme activity inside and outside condensates, we performed two sets of single-turnover decapping experiments using capped radiolabeled RNA and thin-layer chromatography to resolve m^7^GDP product (**Figure 4A**). In the first experiment, we measured decapping in a mixture containing Dcp1/Dcp2_ext_ inside (dense phase) and outside of droplets (dilute phase) by monitoring m^7^GDP formation after addition of capped RNA substrate (bulk decapping). In a second experiment, we separated the dilute and dense phases by centrifugation, added capped substrate to the dilute phase only, and monitored product formation (dilute phase decapping). Any differences observed in decapping activity between the two samples can be attributed to contributions from Dcp1/Dcp2 sequestered in droplets.

**Figure 4.**
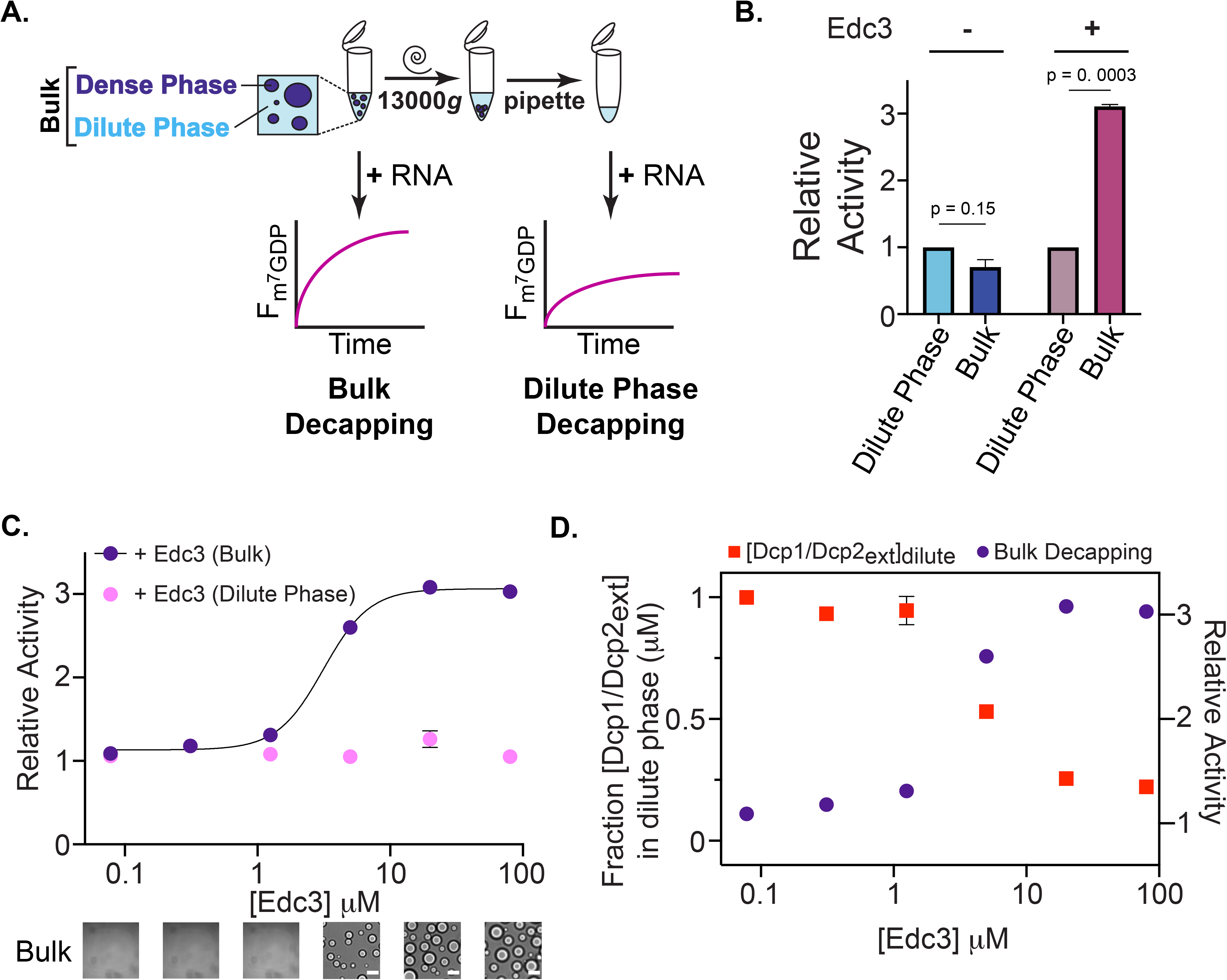
Edc3 activation of decapping is coupled to phase separation. **A**. Schematic of *in vitro* decapping assay to determine contribution of decapping activity from Dcp1/Dcp2_ext_ in droplets (Dense Phase) and outside droplets (Dilute Phase). **B**. Dcp1/Dcp2_ext_ localized in droplets does not contribute substantially to overall activity (blue bars) but removal of Dcp1/Dcp2_ext_/Edc3 droplets strongly diminishes observed activity (pink bars). **C**. Activation of decapping by Edc3 is concomitant with formation of microscopically visible droplets. Removal of droplets abrogates activation by Edc3. **D**. Edc3 activation coincides with depletion of Dcp1/Dcp2_ext_ (red squares) from solution. Curve showing Edc3 activation is reproduced from **C** for purposes of comparison. Error bars represent s.e.m. Scale bar, 10 μm.

We used this approach to determine the contribution of Dcp1/Dcp2_ext_ phase separation to overall activity in the absence or presence of Edc3. We observed comparable rates of decapping in the Dcp1/Dcp2_ext_ dilute and bulk phases, demonstrating Dcp1/Dcp2_ext_ sequestered into droplets does not significantly contribute to the overall activity and confirms Dcp1/Dcp2_ext_ droplets are inactive (**Figure 4B** left blue bars). We do not observe an enhanced repression in the bulk phase using this assay because it also monitors decapping from Dcp1/Dcp2_ext_ remaining outside droplets, which masks the enhanced repression observed in Dcp1/Dcp2_ext_ droplets (**Figure 3**). This is in contrast to droplets containing Edc3 and Dcp1/Dcp2_ext_, whereby activity in the bulk solution is three-fold greater than in the dilute phase, which exhibits activity similar to Dcp1/Dcp2_ext_ (**Figure 4B** right purple bars). Together, these results indicate (i) condensates containing the decapping complex have a weaker enzymatic activity in the absence of Edc3, in agreement with experiments using fluorescently labelled substrate RNA (**Figure 3**), and (ii) condensate composition regulates decapping activity.

To examine this phenomenon more closely, we performed a series of decapping experiments under increasing concentrations of Edc3 and looked for the appearance of phase-separated droplets using microscopy. Cooperative activation of decapping by Edc3 was observed and correlated with phase separation (**Figure 4C**). Multivalency is crucial for this behavior as the cooperativity observed in Edc3 activation was dependent on dimerization (**Figure S4A** and **S4B**). Performing these experiments with only the dilute phase abolished the increase in decapping rate, indicating that decapping in Dcp1/Dcp2_ext_/Edc3 condensates contributes significantly to overall activity (**Figure 4C** pink data points).

We further characterized the ability of Edc3 to recruit Dcp1/Dcp2_ext_ and RNA into droplets using a pelleting assay similar to the decapping assay and fluorescence microscopy. The loss of fluorescently-labeled Dcp1/Dcp2_ext_ from the dilute phase was measured at Edc3 concentrations used in decapping experiments (**Figures 4D, S4C**, and **S4D**). A decrease in Dcp1/Dcp2_ext_ in the dilute phase occurred at Edc3 concentrations coinciding with the formation of phase-separated droplets and Dcp1/Dcp2_ext_ continued to be depleted from solution as Edc3 concentrations were increased (**Figure 4D**). The *K*_1/2_ of Dcp1/Dcp2_ext_ depletion (4.3 μM) is in good agreement with the *K*_1/2_ of Edc3 activation (3.3 μM), indicative of a coupling between activation of decapping and phase separation. Edc3 accumulated in the dilute phase, ruling out its depletion as a mechanism for the differential activity observed in bulk decapping (**Figure S4C**). While variation in Edc3 concentration influenced the partitioning of Dcp1/Dcp2_ext_ in droplets, Edc3 did not significantly affect the partitioning of RNA into droplets, signifying RNA enrichment is not the predominant mechanism for increased decapping we attribute to droplet formation (**Figure S5**). Our results suggest Edc3 reorganizes the protein-protein interactions in condensates and couples catalytic activation to phase separation in order to compartmentalize decapping activity.

### Edc3 shifts a conformational equilibrium in Dcp1/Dcp2 to activate decapping

Dcp1/Dcp2 is a dynamic enzyme that can exist in multiple inactive states thought to precede formation of a catalytically active complex (Mugridge et al., 2018). We used NMR spectroscopy to understand how Edc3 affects Dcp1/Dcp2 dynamics to promote activation of decapping under conditions where Dcp1/Dcp2_ext_ does not phase separate and then asked if a similar mechanism was responsible for decapping activation in condensates. The structured catalytic core of Dcp2 (Dcp2_core_) undergoes large conformational changes in order to accommodate mRNA substrate for cap hydrolysis (**Figure 5A**). In the absence of activators and substrate, Dcp1/Dcp2_core_ exists in an equilibrium between two forms: a highly populated, closed inactive form where the cap binding site and RNA-body binding channel are occluded (closed, inactive) and an open precatalytic form that can bind RNA and is on-pathway to decapping (Wurm et al., 2017). Because interconversion of the precatalytic and inactive forms is fast on the NMR chemical shift timescale, resonance positions report on the population weighted average of these two states in solution. Accordingly, we hypothesized the C-terminus of Dcp2 may influence the equilibrium between precatalytic and inactive forms observable by NMR.

**Figure 5.**
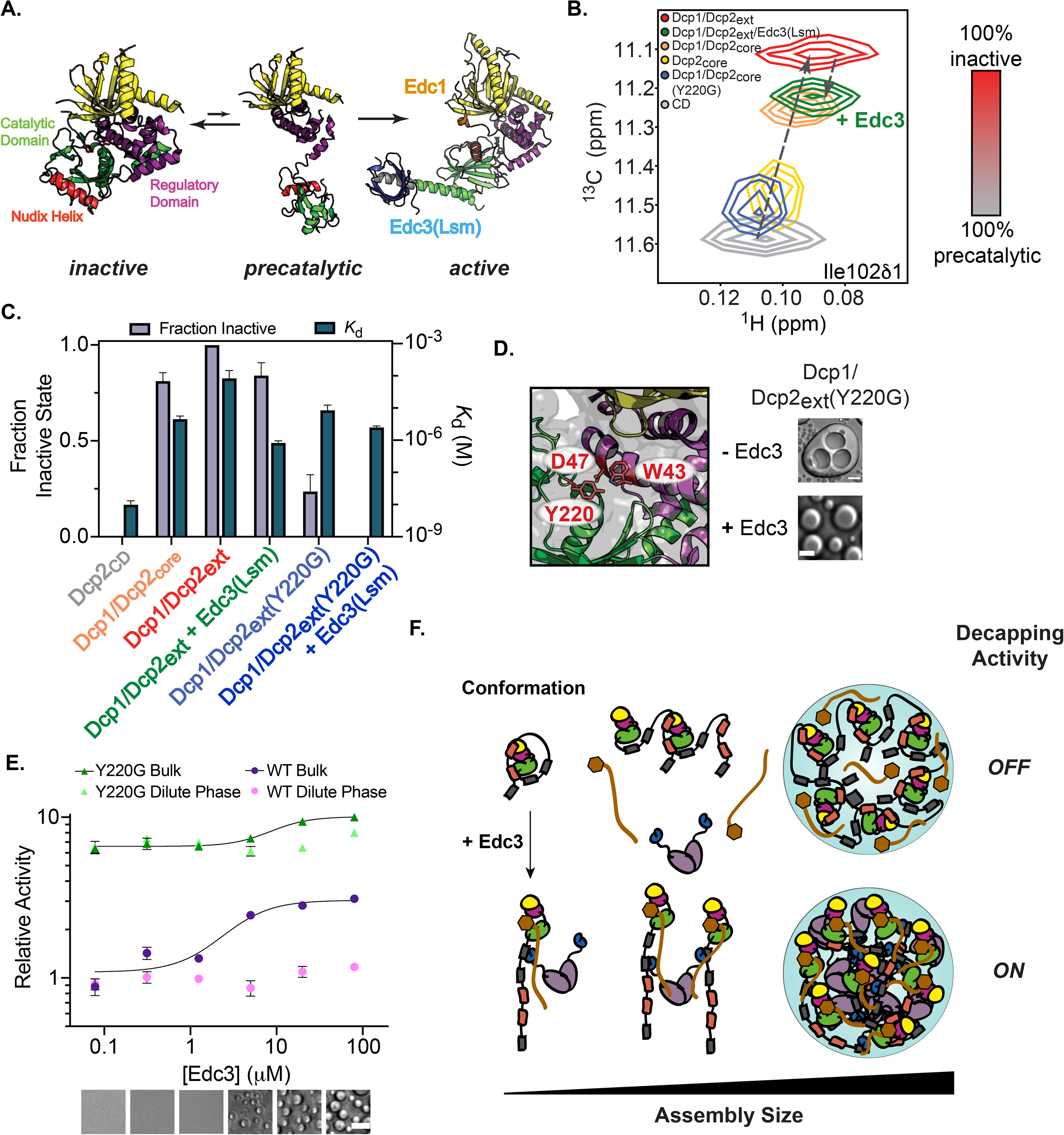
The Dcp2 C-terminus stabilizes an autoinhibited conformation required for regulation of decapping in condensates. **A**. Cartoon representation of the structured core domains of Dcp1/Dcp2 (colored as in Figure 1A). Dcp2 undergoes extensive conformational rearrangements, existing in a fast equilibrium between closed, inactive and open, precatalytic states (PDB: 2QKM) and the addition of activators (Edc1 and Edc3) stabilizes formation of the active state (PDB: 6AM0). The Nudix Helix, which contains the critical catalytic residues is highlighted in red. **B**. ^1^H/^13^C-HSQC of a terminal methyl group in Ile102 undergoes linear chemical shift changes toward a closed, inactive state upon addition of the Dcp2 regulatory domain, Dcp1, and the Dcp2 C-terminus. Edc3 and mutation of Y220 revert the chemical shifts toward the precatalytic state. **C**. Increasing population of the inactive state (gray bars) correlates with weakened RNA binding by Dcp1/Dcp2 (steel blue bars). Edc3 and the Y220G mutation restore RNA binding by destabilizing the inactive state. **D**. (Left) Y220 in the catalytic domain of Dcp2 occludes residues critical for cap recognition (W43 and D47) in the inactive state (PDB: 2QKM). (Right) Droplets formed by Dcp1/Dcp2_ext_(Y220G) do not relax to a spherical state upon fusion (top). Dcp1/Dcp2_ext_(Y220G)/Edc3 droplets have properties similar to wild-type (bottom). Protein concentrations are 150 μM. Scale bar, 10 μm. **E**. The Y220G mutation leads to strong activation of decapping and minimizes differences between activity inside and outside condensates. Brightfield microscopy images correspond to conditions for Dcp1/Dcp2_ext_(Y220G)/Edc3. Purple curves represent kinetics for wild-type Dcp1/Dcp2_ext_/Edc3 droplets. Error bars are s.e.m. Scale bar, 10 μm. **F**. Model showing how Edc3 mediates a conformational change in Dcp1/Dcp2 that is coupled to an alteration of the protein-protein interactions promoting higher-order assemblies found in condensates. These changes in interactions switch decapping activity from an off to on state.

Studying conformational dynamics in proteins containing large IDRs is challenging due to significant overlap of crosspeaks in the NMR spectra caused by the poorly structured IDR, which makes analysis difficult. To overcome this in studying Dcp1/Dcp2_ext_, we prepared a segmentally-labelled Dcp1/Dcp2_ext_ that retains wild-type activity and gives a well-dispersed spectrum, which allows for transfer of assignments from Dcp1/Dcp2_core_ (**Figures S6** and **S7**) (Refaei et al., 2011). We confirmed the aforementioned preexisting equilibrium by showing the regulatory domain of Dcp2 and Dcp1 promote co-linear chemical shift perturbations in several resonances of the catalytic domain that report on the precatalytic and inactive forms (**Figures 5B** and **S8**). We observed the C-terminus causes many of these resonances to undergo additional chemical shift perturbations (**Figures 5B** and **S8** red resonances). Because these perturbations are co-linear and in the direction of the inactive state, we conclude the C-terminus increases the population of the inactive form of Dcp2 (**Figure 5C**). Moreover, we observed numerous chemical shift perturbations in Dcp2 that suggest an interaction or reorganization of the structured domains in the presence of the disordered C-terminus (**Figure S9**). Additionally, the C-terminus disfavors the interaction between Dcp1/Dcp2_ext_ and RNA as shown by a ten-fold increase in the *K*_d_ for RNA relative to the structured core domains (Dcp1/Dcp2_core_) (**Figures 5C** and **S10A**). This is consistent with previous reports showing residues in the catalytic domain of Dcp2 important for interaction with RNA are blocked in the inactive state and, as a result, the population of the inactive state directly correlates with a decrease in binding affinity for RNA (increase in *K*_d_) (Deshmukh et al., 2008; She et al., 2008; Wurm et al., 2017). Together, these results implicate stabilization of the inactive state as the mechanism for autoinhibition and describe why Dcp1/Dcp2_ext_ droplets exhibit minimal decapping.

We next asked if Edc3 alleviates autoinhibition by shifting the conformational equilibrium of Dcp1/Dcp2_ext_ into the precatalytic form that binds RNA more tightly. In order to prevent NMR resonance broadening that may result from phase separation, we assayed activation using the Edc3 Lsm domain, Edc3(Lsm), which is necessary and sufficient for enhancing Dcp2 catalysis and abrogates phase separation (**Figure 2C**) (Paquette et al., 2018). Addition of Edc3(Lsm) to Dcp1/Dcp2_ext_ led to a migration of resonances along the linear trajectory toward the chemical shift for the open conformation, suggesting an increase in the fraction of precatalytic Dcp1/Dcp2_ext_ complex (**Figures 5B** and **S8** green resonances). Quantification of the population of the inactive state across several residues demonstrates Edc3(Lsm) causes Dcp2 to occupy this state ~80% of the time, which is most similar to the structured core domains lacking the C-terminus, Dcp1/Dcp2_core_ (**Figure 5C**). This trend was also observed in a more global analysis of chemical shifts in Dcp2 that showed Edc3(Lsm) reduced many of the chemical shift perturbations we observed upon appending the C-terminal extension to the core domain and indicates interactions between the C-terminus and core domains are reduced (**Figure S9**). The addition of Edc3(Lsm) also restores RNA binding by Dcp1/Dcp2_ext_ as the reported *K*_d_ for RNA decreased 100-fold (**Figures 5C** and **S10A**). We conclude Edc3 activates decapping by increasing the amount of precatalytic conformation, which enables tight RNA recognition and subsequent cap hydrolysis, explaining why only droplets containing the three proteins together are capable of performing robust decapping.

### Conformational regulation of Dcp2 is required for compartmentalization of decapping in condensates

In previous work, we identified a conserved aromatic residue in the catalytic domain of Dcp2 (Y220) contacts residues critical for m^7^G recognition (W43 and D47), and predicted this residue stabilizes the inactive state (**Figure 5D**). Consistent with this view, mutation of Y220 alleviates autoinhibition and bypasses Edc3 activation (Paquette et al., 2018). Using NMR, we determined whether this gain-of-function arises from a change in the precatalytic-inactive equilibrium in Dcp1/Dcp2. The chemical shifts for resonances reporting on this equilibrium underwent significant perturbations toward the precatalytic state, indicating this mutation disrupts formation of the inactive conformation (**Figure 5B** and **5C**). In addition, the Y220G mutation in Dcp2 restored RNA binding in Dcp1/Dcp2_ext_, making it insensitive to the presence of Edc3 (**Figures 5C** and **S10A**). These observations confirm Edc3 activates decapping by promoting formation of a precatalytic conformation of Dcp1/Dcp2 that makes high affinity interactions with substrate RNA. The Y220G mutation in Dcp2 bypasses the requirement for Edc3 by also promoting the precatalytic conformation.

Interactions important for autoinhibition are also crucial for Dcp1/Dcp2_ext_ droplet formation (**Figure 2C** and **2D**). We next asked if there is a coupling between phase separation and the conformational equilibrium of Dcp1/Dcp2_ext_ using the Y220G mutation. While droplet formation was not abolished in Dcp1/Dcp2_ext_(Y220G), the droplets became less liquid-like as they failed to properly relax after fusion, suggesting their physicochemical properties are altered (**Figure 5D** top right compared to top row **Figure 1C**) (Alberti et al., 2019). Conversely, Dcp1/Dcp2_ext_(Y220G)/Edc3 droplets presented physicochemical properties similar to wild-type (**Figure 5D** bottom right compared to bottom row **Figure 1C**). This suggests the conformational equilibrium of Dcp1/Dcp2 may be coupled to the material properties of condensates.

The Y220G mutation in Dcp2 bypasses activation by destabilizing the inactive state of Dcp1/Dcp2 (**Figure 5B** and **5C**). If Edc3 is no longer able to conformationally regulate Dcp1/Dcp2, then its ability to couple activation and phase separation may be impaired. Decapping assays performed on Dcp1/Dcp2_ext_ containing the Y220G mutation revealed 7-fold activation in basal decapping (**Figure 5E**). Dose-dependent addition of Edc3 resulted in phase separation similar to wild-type, however the extent of activation by Edc3 was minimized, only activating decapping by 1.5-fold compared to 3-fold in wild-type (**Figure 5E**). In addition, the *K*_1/2_ of activation is 3-fold greater than for wild-type and no longer corresponds well with phase separation (**Figure S10B**). Thus, the Y220G mutant has a dual effect on decapping: (i) it accelerates decapping to the extent activity inside or outside condensates is indistinguishable and (ii) it decouples Edc3-mediated activation from phase separation. From these results, we conclude the conformation of Dcp2 is crucial for proper regulation of activity and compartmentalization of mRNA decapping.

## DISCUSSION

We addressed the importance and mechanisms of phase separation in RNA decapping by Dcp1/Dcp2. We found decapping can be both repressed and activated in condensates depending on condensate composition. We explain this differential activity in condensates by demonstrating changes in Dcp1/Dcp2 conformation are coupled to alterations in the protein network underlying condensate formation. IMs in Dcp2 stabilize an autoinhibited conformation and drive self-association into condensates repressed in decapping activity (**Figures 2, 3, 4, 5**). The activator Edc3 rectifies this inhibited environment by interacting with HLMs in Dcp2, to cause a conformational change in Dcp1/Dcp2 important for substrate recognition and subsequent catalysis (**Figures 2** and **5**). These Edc3-mediated interactions rewire condensates to promote mRNA decapping (**Figures 3** and **4**). Our work provides the most direct evidence that highly regulated decapping can occur in phase-separated condensates, and suggests composition couples meso- and Ångstrom-scale organization of enzyme active sites to modulate activity and control the function of condensates. Further, as discussed below the specific nature of the regulation suggests new roles for condensates in RNA biology.

We find an emergent property of phase separation is the ability to amplify activation of enzymes through the dense network of protein-protein interactions. Edc3 activates decapping by as much as 80-fold in droplets, which is in contrast to the three-fold stimulation determined in bulk experiments (**Figures 3** and **4**). The increased activation observed in droplets is not due to an acceleration of the rate of decapping but rather an inhibition of Dcp1/Dcp2 activity in the condensate environment (**Figure S4**). Therefore, we propose activation of decapping in condensates is greater than in the surrounding solution because condensates strongly repress decapping activity.

How might repression of enzyme activity be greater inside condensates? We show that inhibitory motifs in the intrinsically disordered C-terminus of Dcp2 can shift a conformational equilibrium in the active site to favor an autoinhibited conformation (**Figure 5B** and **5C**). It is possible this equilibrium is shifted further in condensates to repress enzyme activity. Consistent with this view, a mutation in the active site of Dcp2 (Y220G) shifts the conformational equilibrium to alleviate autoinhibition and alters the material properties of Dcp1/Dcp2 condensates (**Figure 5**). These observations suggest active site conformation is coupled to long-range interactions that occur between Dcp1/Dcp2 molecules in condensates to directly regulate activity. Previous work demonstrated how conformational changes are important for promoting phase separation of heterochromatin and our results suggest phase separation has the ability to influence atomic fluctuations in resident proteins, leading to potential new mechanisms of regulation (Sanulli et al., 2019). How active site conformation is coupled to mesoscale organization in condensates is an exciting area of research that is a challenge for future work.

Activation of decapping in condensates is promoted by Edc3, which binds HLMs in the C-terminus of Dcp2 to cause changes in both the molecular interactions driving phase separation and the conformation of the decapping complex (**Figures 2, 3**, and **5**). In addition, the multivalent interactions between Edc3 and Dcp2 couple phase separation and activation of decapping to compartmentalize decapping (**Figure 4**). Analogous to the Y220G mutation, Edc3 promotes the open conformation of Dcp1/Dcp2 to increase its interaction with RNA 100-fold (**Figure 5B** and **5C**). These data indicate formation of the open state of Dcp1/Dcp2 is a critical step in activation of decapping.

Macromolecular composition is important for regulating specificity in biochemical reactions within condensates (Banani et al., 2016). We propose composition serves as a mechanism to directly inhibit or activate enzymatic activity in condensates. In the framework of decapping, cells may leverage this feature to minimize aberrant mRNA decapping events that would result from widely distributed decapping mRNPs and buffer against changes in cellular environment. In support of this, we observed decapping rates remained constant inside and outside condensates once phase separation occurred despite significant changes in Dcp1/Dcp2_ext_ and Edc3 concentrations (**Figures 4, 4D**, and **S4**). Our data suggest Edc3 increases the partitioning of Dcp1/Dcp2 into condensates while Edc3 partitioning is reduced to activate residual Dcp1/Dcp2 outside condensates (**Figure S4C** and **S5A**). In this way the thermodynamic coupling of Dcp1/Dcp2_ext_ and Edc3 phase separation leads to a buffering of decapping activity. A buffering mechanism has been posited for the control of gene expression through condensation of transcriptional regulators and demonstrates how cells utilize phase separation to regulate gene expression from transcription to degradation (Hnisz et al., 2017). The Y220G mutation decouples activation from phase separation to cause rapid decapping with rates that are indistinguishable inside or outside condensates. The widespread, accelerated decapping could explain the temperature sensitive phenotype observed for the Y220 mutation in budding yeast (Deshmukh et al., 2008). Furthermore, the regulation of decapping by phase separation could explain why removal of the Dcp2 C-terminus leads to dysregulation in mRNA abundance in yeast, as the decapping protein is now unleashed to promiscuously target atypical decapping substrates and promote their degradation (He et al., 2018).

Our work on Edc3 has implications for other activators of decapping that interact with intrinsically disordered regions in the decapping complex (Jonas and Izaurralde, 2013). For example, like Edc3, the conserved mRNA decay factor Pat1 interacts with HLMs to activate decapping (Charenton et al., 2017; Lobel et al., 2019). Thus, we predict Pat1 and similar cofactors stimulate decapping by altering the conformation of the Dcp2 active site. While these factors may utilize a common mechanism to activate decapping, they regulate distinct subsets of mRNAs (Chang et al., 2007; He et al., 2018). Pat1 and other HLM-binding proteins are enriched in P-bodies and could serve to alter the specificity and activity of Dcp2 on mRNAs recruited to cellular condensates (Xing et al., 2018).

How can P-bodies serve as both sites of mRNA storage and decay in cells? Our investigation of decapping in condensates using a minimal *in vitro* system suggests the molecular composition of P-bodies could be regulated to determine their functional state depending on cellular context. Furthermore, the direct observation of m^7^G cap status in condensates using a dual-labeled RNA substrate revealed that cleavage of the cap does not lead to loss of mRNA from condensates (**Figure 3**). The retention of an intact RNA body within the phase-separated compartment raises the possibility transcripts within a P-body can be decapped but not degraded. This stored RNA transcript could then be released from a P-body for subsequent decay in the cytoplasm or be recapped and re-enter the translating pool (Trotman and Schoenberg, 2019). Thus, a P-body can serve as an additional, dynamic cellular checkpoint in RNA degradation, providing a mechanism to ensure or prevent a particular fate for a transcript.

Different from our observations, previous *in vitro* analysis of Dcp1/Dcp2/Edc3 sequestered in phase-separated droplets indicated decapping activity was inhibited (Schütz et al., 2017). Excess RNA was used in the initial study while we monitored decapping under single-turnover experiments where protein concentrations exceed substrate RNA concentrations. Taken together, these results suggest RNA concentration may provide an additional mechanism for regulation of decapping activity in condensates. More broadly, RNA often has a critical role in the assembly of condensates and can regulate the properties and control the specificity of interactions in condensates (Van Treeck and Parker, 2018). Thus, characterizing the relationship between substrate and enzymatic activity in condensates will likely be important for understanding how phase separation regulates biochemical reactions.

The observation that catalytically active Dcp1/Dcp2_ext_/Edc3 droplets were often smaller and more homogenous in size provides *in vitro* evidence that small, microscopically invisible P-bodies may be the relevant sites for 5’-3’ mRNA decay (**Figure S1E**). This is in support of recent computational modeling that demonstrated small, numerous P-bodies are most efficient to degrade cellular transcripts (Pitchiaya et al., 2019). These smaller mRNPs may have a more discrete composition than larger membraneless organelles, which instead represent a broader collection of decay factors and mRNAs sequestered for storage. While finding support of our observation *in vivo* requires improved visualization methods, ultra-high resolution microscopy and nanoparticle tracking have emerged as a tractable method for visualizing diffraction-limited P-bodies (Rao and Parker, 2017). Regardless, the organization of 5’-decay factors in liquid-like mRNPs represents a highly dynamic means to regulate RNA processing.

Phase separation has been shown to be important for the regulation of numerous enzymatic processes involving mRNA in cells, including translation, deadenylation, and miRNA processing (Hondele et al., 2019; Kim et al., 2019; Sheu-Gruttadauria and MacRae, 2018). The fluorescent probes developed here can be used to examine molecular mechanisms of other condensates important for RNA biology. We demonstrate the importance of mesoscale phase-separated assemblies in regulating mRNA decapping. The composition-dependent activation of decapping in condensates may be utilized by cells to regulate whether P-bodies are sites of mRNA decay or storage. The interplay between composition, molecular interactions, and enzyme conformation in Dcp2 condensates underscore the complexity of phase separation in cellular processes. Emergent properties equip cells with highly regulatable sites of enzyme activity that support multiple biological outcomes.

## ACKNOWLEDGEMENTS

We thank Kari Herrington and the Center for Advanced Light Microscopy at UCSF for guidance and technical assistance in collecting microscopy data. We also thank Marcin Warminski and Agnieszka Mlynarska-Cieslak (UW) for providing reagents for RNA labelling and Mark Kelly for assistance through the UCSF NMR Facility. The authors thank Christian Freund, Aashish Manglik, and members of the J.D.G. lab for experimental guidance and many helpful discussions. We extend thanks to Geeta Narlikar and Stephen Floor for useful discussions in writing the manuscript. This work was supported by US National Institutes of Health (R01 GM078360 to J.D.G.), Foundation for Polish Science (TEAM/2016-2/13 to J.J.), National Science Centre, Poland (UMO-2018/31/B/ST5/03821 to J.K.), a Moritz-Heyman UCSF Discovery Fellowship (to R.W.T.), and an ARCS Foundation Fellowship (to R.W.T.).

## AUTHOR CONTRIBUTIONS

All authors designed research plan. J.D.G. and R.W.T. designed experiments and A.D., J.K., and J.J. designed and synthesized the dual-labelled RNA probe. R.W.T. purified proteins and performed experiments. All authors contributed to the writing and editing of the manuscript. J.K., J.J., and J.D.G supervised research.

## DECLARATION OF INTERESTS

The authors declare no competing interests.

## METHODS

### Protein Expression and Purification

The *S*. *pombe* Dcp1(1-127) and Dcp2_ext_(1-504) DNA sequences obtained from Integrated DNA Technologies were codon-optimized for *E. coli* and cloned into MCS1 and MCS2 of a pRSF co-expression vector, respectively. A N-terminal His_6_-MBP-TEV was cloned into the vector upstream of Dcp1 and Dcp2 constructs except 96-243, 1-243, 1-266, and 1-266_eSrtA_ also contained a StrepII tag at the C-terminus. All truncation constructs of Dcp2 and point mutants were cloned using divergent primers and whole plasmid PCR from this initial vector (See Supplemental Table 1 for constructs used in this study).

Protein constructs were expressed in *E. coli* BL21(DE3) (New England Biolabs) grown in LB medium. Cells were grown at 37°C until they reached an OD_600_ = 0.6–0.8 and then transferred to 4°C for 30 min before induction by addition of 0.75 mM IPTG (final concentration). Cells were induced for 16–18 hours at 20°C. Cells were harvested at 5000*g*, resuspended in lysis buffer (25 mM HEPES pH 7.5, 400 mM NaCl, 10 mM 2-mercaptoethanol, 0.1% Triton X-100 supplemented with lysozyme and Roche cOmplete protease inhibitor tablet), lysed by sonication (50% duty cycle, 4 × 1 min), and clarified at 16,000*g*. Proteins were purified in a two-step affinity purification: GE Healthcare StrepTrap affinity column followed by a GE Healthcare HiTrap Heparin column. Briefly, proteins were loaded onto StrepTrap and then washed with 10 column volumes (CV) lysis buffer without detergent followed by a second wash with 10 CV 25 mM HEPES pH 7.5, 100 mM NaCl, 10 mM 2-mercaptoethanol. Elution of target proteins was performed using a step elution to 25 mM HEPES pH 7.5, 100 mM NaCl, 1 mM DTT, 5 mM desthiobiotin. This elution was directly loaded onto a HiTrap Heparin column equilibrated with low salt buffer (25 mM HEPES pH 7.5, 100 mM NaCl, 1 mM DTT). 10CV of the low salt buffer was used to wash the column and proteins were eluted over a 10CV gradient to 100% high salt buffer (25 mM HEPES pH 7.5, 1 M NaCl, 1 mM DTT). For Dcp1/Dcp2 constructs lacking a C-terminal StrepII tag, a GE Healthcare HisTrap column was used in place of the StrepTrap. The lysis buffer above was supplemented with 10 mM imidazole and the column was washed with 10CV lysis buffer without detergent and 10CV 25 mM HEPES pH 7.5, 100 mM NaCl, 10 mM imidazole, 10 mM 2-mercaptoethanol. Proteins were eluted from the HisTrap column in 10CV 25 mM HEPES pH 7.5, 100 mM NaCl, 250 mM imidazole, 10 mM 2-mercaptoethanol and incubated with TEV overnight. The TEV digested proteins were applied to a HiTrap Heaprin column as described above. All protein constructs were further purified by size-exclusion chromatography on a GE Healthcare Superdex 200 16/60 column equilibrated with a final buffer (25 mM HEPES pH 7.5, 150 mM NaCl, 1 mM DTT). Proteins were then concentrated, flash frozen in liquid nitrogen and stored at −80°C.

A His_6_-TEV-Edc3 plasmid was generated in a pET30b plasmid, which already contained a His_6_-TEV coding sequence. The Edc3(1-94) construct containing the N-terminal Lsm domain was created by whole-plasmid PCR with 5’-phosphorylated primers. The Edc3 constructs were purified by Nickel affinity chromatography as described above followed by TEV digestion overnight at room temperature. Cleaved Edc3 was further purified by size-exclusion chromatography as described for Dcp1/Dcp2.

A pET29 plasmid encoding a pentamutant SortaseA(Δ59) construct from *S. aureus* with improved activity (eSrtA) was obtained from Addgene (plasmid #75144) and expressed in LB medium at 18°C for 16 hours (Chen et al., 2011). Cells were pelleted at 5000*g* and resuspended in lysis buffer containing 50 mM Tris pH 8, 300 mM NaCl, 10 mM imidazole, and 1 mM MgCl_2_ supplemented with a Roche cOmplete protease inhibitor tablet. Following sonication (50% duty cycle, 4 ’ 1 min) the lysate was clarified at 16000*g* for 30 minutes at 4°C. Clairifed lysate was then passed over a GE HisTrap Nickel affinity purification column. eSrtA was eluted in 25 mM HEPES pH 7.5, 150 mM NaCl, 250 mM imidazole, dialyzed against 25 mM HEPES pH 7.5, 150 mM NaCl overnight at 4°C, concentrated to 1 mM final concentration as determined by A_280_, and flash frozen in liquid nitrogen for storage at −80°C.

### Fluorescent-labeling of purified proteins and RNAs

Fluorescent-labeled Dcp1/Dcp2_ext_ and Edc3 were generated by diluting the protein to 0.5 mg/mL in size exclusion buffer without DTT and dialyzing in this buffer at 4°C for at least four hours. 5-fold molar excess Cy5 maleimide or Fluorescein maleimide (Thermo-Fisher) were added to the protein solution and incubated for one hour at room temperature. The reaction was quenched by addition of β-ME to a final concentration of 10 mM. Free dye was separated from labelled protein by Illustra NICK columns (GE Healthcare) according to the manufacturer’s instruction. The protein was then exchanged back into size exclusion buffer containing DTT by concentrating and diluting 10-fold three times. Labelling efficiency and concentrations were calculated by UV-Vis spectroscopy. Labelled oligonucleotides for fluorescence polarization and RNA enrichment experiments were purchased from Integrated DNA Technologies with a 5’ 6-FAM modification.

### Brightfield and Fluorescence Microscopy

Microscopy images were collected on an inverted widefield fluorescence Nikon Ti-E microscope equipped with a Hamamatsu Flash4.0 camera using either PlanApo 20x or 40x air objectives. Samples were imaged in a Greiner Bio-One 384-well glass bottom plate PEGylated using 20 mg/mL PEG-Silane (Laysan Bio, MPEG-SIL-5000) and passivated with 100 mg/mL BSA as described (Keenen et al., 2018). Prior to addition of samples, the wells were washed 3x with 25 mM HEPES pH 7.5, 150 mM NaCl, 1 mM DTT. Dcp1/Dcp2 constructs assayed by microscopy for phase separation were prepared by initiating removal of the N-terminal MBP solubility tag with 1:40 molar equivalent TEV:Dcp1/Dcp2. Dcp1/Dcp2/Edc3 droplets were prepared by incubating Dcp1/Dcp2 and Edc3 prior to removal of the N-terminal MBP tag from Dcp1/Dcp2_ext_. TEV cleavage and droplet settling were allowed to occur for 30 minutes at room temperature prior to imaging. For localization and enrichment of Dcp1/Dcp2_ext_, Edc3, or RNA in droplets, samples were prepared so that 5% of a given component was fluorescently labelled. Image analysis was performed in FIJI (Schindelin et al., 2012).

### Synthesis of dually labelled RNA probe

The reagents for RNA labelling, FAM-m^7^Gp_3_A_m_pG and pAp-SCy5, were synthesized by modifications of previously reported methods (Kowalska et al., 2012; Warminski et al., 2017). The detailed synthetic procedures will be published elsewhere. FAM-m^7^G-capped RNA was generated on the template of annealed oligonucleotides, which contained a T7 Aϕ2.5 promoter sequence (CAGTAATACGACTCACTATT) and encoded a 35-nt-long sequence (AGG GAAGCG GGCATG CGGCCA GCCATA GCCGAT CA), (the first and second transcribed nucleotides are marked in red and green, respectively). Typical *in vitro* transcription reaction (100 μL) was carried out at 37 °C for 4 h and contained RNA Pol buffer (40 mM Tris-HCl pH 7.9, 6 mM MgCl_2_, 1 mM DTT, 2 mM spermidine), 10 U/μL T7 polymerase (ThermoFisher Scientific), 1 U/μL RiboLock RNase Inhibitor (ThermoFisher Scientific), 0.5 mM CTP/GTP/UTP, 0.125 mM ATP, 0.625 mM FAM-m^7^Gp_3_A_m_pG cap and 0.1 μM annealed oligonucleotides as a template. Following 4 h incubation, the template was removed by treatment with 1 U/μL DNase I (ThermoFisher Scientific) for 30 min at 37 °C. The crude RNAs were purified using RNA Clean & Concentrator-5 (Zymo Research). Transcripts quality was checked on 15% acrylamide/7 M urea gels, whereas the concentration was determined spectrophotometrically.

The obtained transcripts were directly used in the ligation step with a Sulfo-Cy5 (SCy5) labelled pAp analogue. A typical ligation reaction (30 μL) was carried out at 16 °C overnight and contained 5’ capped RNA (1 μM), 1 U/μL T4 RNA ligase 1 (New England Biolabs), 1.3 U/μL Ribolock RNase inhibitor (ThermoFisher Scientific), 100 μM pAp-SCy5 analogue, 0.1 volumes of DMSO (3 μL), 0.03 volumes of 0.1 M DTT (1 μL) and 0.1 volumes of 10 mM ATP (3 μL). The resulting dually labelled RNA was first purified using RNA Clean & Concentrator-5 (Zymo Research) followed by the final HPLC purification (Clarity^®^ 3 μM Oligo-RP phenomenex column, linear gradient from 5% to 35% ACN in 50 mM TEAAc pH 7 over 15 min at 50 °C, Agilent Technologies Series 1200 HPLC). The collected fractions were freeze dried 3 times. RNA quality, before and after each purification step, was checked on 15% acrylamide/7 M urea gels (Figure S3), whereas the concentration was determined spectrophotometrically.

### Visualization of Decapping by Microscopy

Dcp1/Dcp2_ext_ droplets were prepared at 60 μM in 25 mM HEPES pH 7.5, 150 mM NaCl, 1 mM DTT supplemented with RNase inhibitor as described above. For reactions containing Dcp1/Dcp2_ext_/Edc3, the decapping complex was monitored at 5 μM and the concentration of Edc3 was 80 μM. The dual-labeled RNA probe was added at a final concentration of 100 nM to both reactions. Initial images were taken to observe localization within droplets. Decapping was initiated by addition of MgCl_2_ to a final concentration of 5 mM while a control reaction was performed without addition of metal. For experiments visualizing activation at various concentrations of Edc3, Dcp1/Dcp2_ext_ droplets were preformed in the presence of 100 nM probe by incubation with TEV for 30 minutes at room temperature. Edc3 was added 10 minutes prior to initiation of the reaction with 5 mM MgCl_2_. Images in both the fluorescein and Cy5 channels were collected at specified time points in Figure 4. The mean droplet intensity at each time point for both channels was background corrected and calculated in FIJI. Mean intensity was plotted as function of time and error bars represent the standard error of the mean for at least ten individual droplets monitored over the course of an experiment. The best fit line representing loss of enrichment of FAM-m^7^GDP from droplets is either a linear function (Dcp1/Dcp2_ext_) or first-order exponential decay function (droplets containing Edc3).

### In Vitro Decapping Assays

For all decapping assays, synthetic 5’-triphosphate 29mer derived from the MFA2 gene of *S. cerevisiae* was purchased from TriLink BioTechnologies and enzymatically capped with GTP[α-^32^P] and S-adenosylmethionine (SAM) as previously described (Jones et al, 2008). Reactions were carried out in 25 mM HEPES pH 7.5, 150 mM NaCl, 5 mM MgCl_2_, 0.1 mg/mL acetylated BSA with RNase inhibitor. Prior to performing assays monitoring Edc3 activation, the MBP tag was removed from Dcp1/Dcp2_ext_ using TEV and separated using Amylose resin. For reactions monitoring Edc3 activation, Dcp1/Dcp2_ext_ was kept constant at 5 μM while Edc3 was varied from 78.1 nM to 80 μM. Dcp1/Dcp2_ext_ concentrations were always at least 10-fold excess over RNA to prevent product inhibition. For reactions examining contribution of only phase-separated Dcp1/Dcp2_ext_ the concentration was increased to 100 μM. 1.5x protein and 3x RNA solutions were equilibrated for 30 minutes separately at room temperature to allow formation of liquid droplets in the protein solution. In reactions examining activity of Dcp1/Dcp2_ext_ remaining outside of liquid droplets, following the 30 minute incubation the protein solution was centrifuged for 10 minutes at 13000*g* to pellet droplets and the supernatant was carefully removed for subsequent assays. Reactions were initiated by mixing 15 μL 1.5X protein and 7.5 μL 3X RNA solutions. Time points were taken by quenching the reaction with excess EDTA and TLC was used to resolve product m^7^GDP from RNA. The formation of product was quantified using a GE Healthcare Typhoon scanner and ImageQuant software and observed rates, *k*_obs_ were determined by fitting to a first-order exponential. Relative rates were determined by normalizing the observed rates to that obtained for 5 μM Dcp1/Dcp2 in the absence of Edc3. In the case of examining activation by Edc3, *k*_obs_ versus Edc3 concentration was fit to the model:

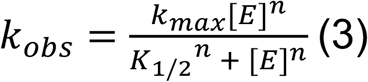

in order to obtain *k*_max_, *K*_1/2_, and *n*. Errors reported for *k*_obs_ and Relative Activity represent the standard error of the mean for at least two independent replicates and errors in *K*_1/2_, *k*_max_, and *n* are the standard error from the obtained fits to equation (3). See Supplemental Tables 2-4 for relative activity values obtained in this study.

### Dcp1/Dcp2_ext_ precipitation assay

Loss of Dcp1/Dcp2_ext_ from solution as a result of sequestration in phase-separated liquid droplets as a function of Edc3 concentration was monitored by loss of fluorescent Dcp1/Dcp2_ext_ signal. Briefly, 5 μM Dcp1/Dcp2 containing 5% labeled with Cy5 maleimide was incubated with Edc3 concentrations between 78.1 nM and 80 μM as in the *in vitro* decapping assay experiments. Following a 30 minute incubation at room temperature, droplets were pelleted by centrifugation at 13000*g* for 10 minutes. The supernatant was carefully removed and placed in a Greiner Bio-One 384-well low volume low binding plate. Fluorescence intensity readings were taken on a BioTek Synergy H4 plate reader and normalized to give a fractional amount remaining in solution. The reported results represent the mean and standard error for three independent experiments. In addition, the supernatant was analyzed using denaturing 4-12% Tris-glycine SDS-PAGE and imaged using a BioRad ChemiDoc. Disappearance of Dcp2 in the lanes was quantified using a region of interest of the same size in FIJI and background subtracting. Only loss of Dcp2 was monitored given the ability to monitor its depletion by both Instant Blue staining and in-gel fluorescence.

### Expression of labeled Dcp1/Dcp2 for NMR

ILVMA methyl labeling of Dcp2 or Dcp1/Dcp2 constructs was carried out in D2O M9 minimal media with ^15^NH_4_Cl and ^2^H /^12^C-glucose as the sole nitrogen and carbon sources, respectively, and labeled precursors (Ile: 50 mg L^−1^, Leu/Val: 100 mg L^−1^, Met: 100 mg L^−1^, Ala: 100 mg L^−1^) were added 40 minutes prior to induction. Following overnight induction with 1 mM IPTG at 20-25°C, cells were lysed by sonication and clarified at 16000*g* as described above. Labeled Dcp2 constructs were purified using a GE Healthcare HisTrap as described above. The His_6_-MBP-TEV tag was removed by overnight digestion with TEV protease at room temperature. Cleaved protein was isolated by performing Heparin affinity chromatography as above and a final purification step using Superdex 200 16/60 size exclusion chromatography equilibrated in 25mM HEPES pH 7.5, 150 mM NaCl, 1 mM DTT.

### SortaseA Ligation of Dcp1/Dcp2 for NMR

Dcp1/Dcp2(1-266) containing a C-terminal LPETGGH *S. aureus* SortaseA recognition site Dcp1/Dcp2(1-266)_eSrtA_ was expressed in a perdeuterated background and ^1^H/^13^C-labeled at ILVMA terminal methyl groups and purified as described above. Purified protein was then mixed with at least five-fold molar excess His_6_-MBP-G_3_-Dcp2(274-504)-StrepII expressed in LB medium and purified as described above. An amount of eSrtA enzyme equal to 0.5 molar equivalent of Dcp1/Dcp2(1-266)_eSrtA_ and 2 mM CaCl_2_ were added to the solution prior to initiation of the reaction with TEV protease, which cleaves the His_6_-MBP tag from G_3_-Dcp2(274-504)-StrepII to expose the required N-terminal glycine for ligation. The reaction was dialyzed against 25mM HEPES pH 7.5, 150 mM NaCl, 2 mM CaCl_2_ overnight at 4°C. Ligated Dcp1/Dcp2(1-504)_eSrtA_ was purified by GE Heparin and GE StrepTrap affinity chromatography as described above. A final dialysis in 25 mM HEPES pH 7.5, 150 mM NaCl, 1 mM DTT was performed overnight at 4°C prior to concentration for NMR experiments.

### NMR Experiments

Prior to data acquisition, NMR samples were exchanged into a buffered D2O solution containing 25 mM HEPES pD 7.1, 150 mM NaCl, and 1mM DTT using either dialysis or a series of concentration and dilution steps in an Amicon centrifugal concentrator. All NMR experiments were performed on an 800 MHz Bruker Avance III or Avance NEO spectrometer equipped with a cryoprobe. Spectra were recorded at 298K, processed using NMRPipe, visualized and analyzed using NMRFAM-Sparky (Lee et al., 2015). Note the concentration of ligated Dcp1/Dcp2_ext_ was kept below the critical concentration in order to prevent confounding effects from phase separation. Experiments examining the effect of Edc3 utilized the Lsm domain (1-94) to prevent phase separation. For the calculation of the inactive state population (p*_inactive_*) of the various Dcp2 constructs, we focused on resonances undergoing linear perturbations indicative of fast exchange. The observed chemical shift was taken as the weighted average of the resonances corresponding to the open and inactive state: the catalytic domain of Dcp2 and Dcp1/Dcp2_ext_, respectively:

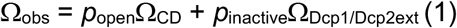

Where Ω_obs_ is the observed chemical shift, Ω_CD_ is the chemical shift of the catalytic domain, and Ω_Dcp1/Dcp2(1-504)_ is the chemical shift corresponding of the eSrtA-ligated construct (all in ppm).

From this relationship, the population of the inactive state can be calculated as:

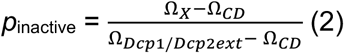

where Ω_X_ is the chemical shift for a given construct of Dcp1/Dcp2 (in ppm). The reported *p*_inactive_ represents the average of four resonances where all constructs were observable.

### Fluorescence Polarization

Fluorescence polarization was performed in Greiner Bio-One 384-well low volume, low binding plates and the final condition for all binding assays was 25 mM HEPES pH 7.5, 100 mM NaCl, 5 mM MgCl_2_, 0.02% Triton X-100, 0.1 mg/mL acetylated BSA and 4U RNase inhibitor. 5’-phosphorylated oligo 30U RNA with 3’ 6-FAM was ordered from Integrated DNA Technologies and used at a final concentration of 5 nM. Reactions were performed in triplicate and incubated for ten minutes before measuring polarization on a LJL Biosystems Analyst AD plate reader. Equilibrium dissociation constants (K_d_) were fit to the Hill equation for single-site binding:

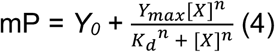

where [X] represents the concentration of protein, Y_0_ is the mP for the probe alone, Y_max_ is the mP value at saturation, and *n* is the Hill coefficient. K_d_ values reported represent the mean and standard error from two independent replicates. Note that for polarization experiments using Dcp1/Dcp2_ext_, the MBP tag was retained to prevent scattering effects from phase separation. The effect of Edc3 on RNA binding was carried out using a construct of Edc3 only containing the Lsm domain (1-94) to prevent scattering effects from phase separation.

---

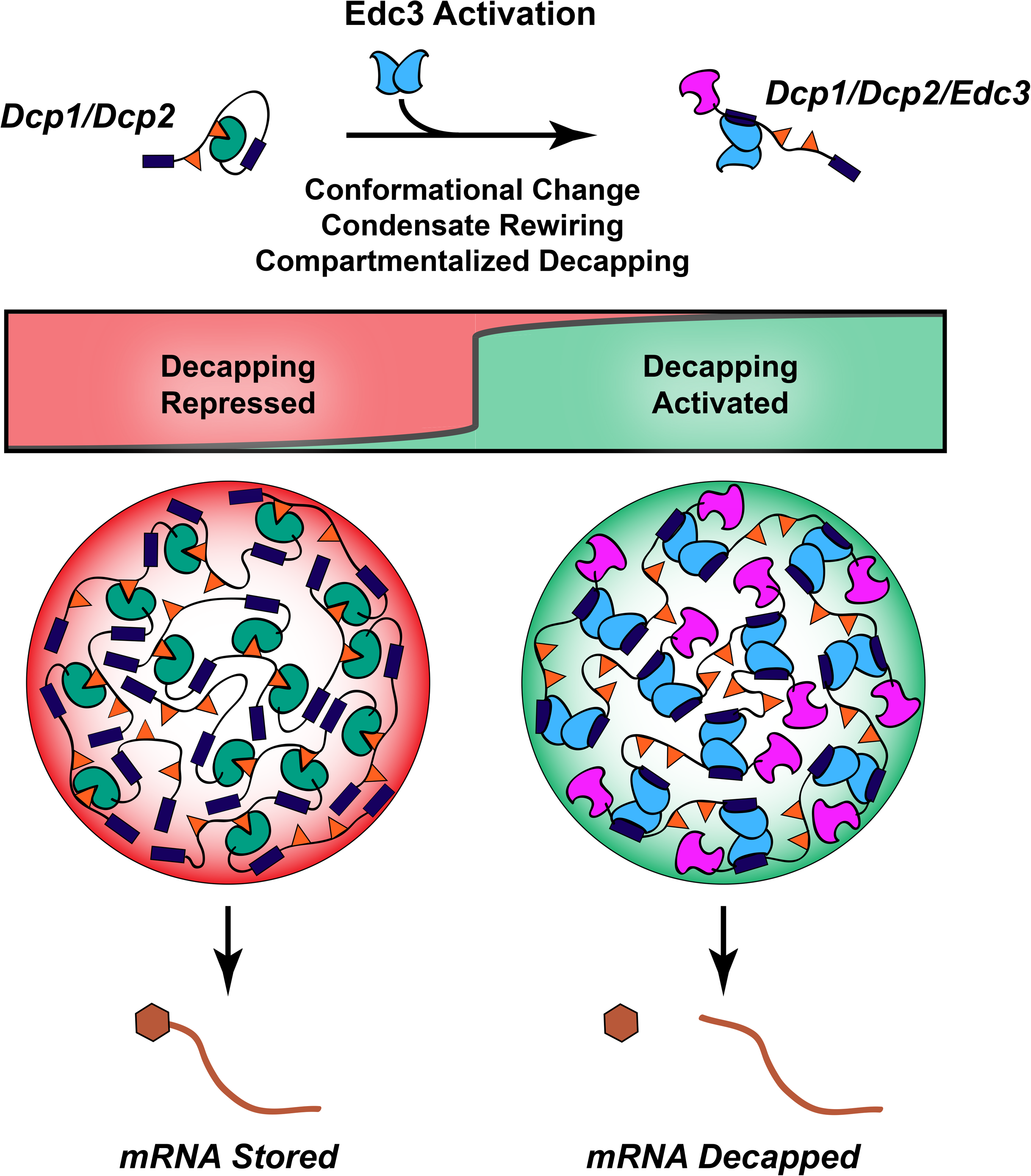

## Notes

### Competing Interest Statement

The authors have declared no competing interest.

## REFERENCES

Abernathy, E., Glaunsinger, B., 2015. Emerging roles for RNA degradation in viral replication and antiviral defense. Virology, 60th Anniversary Issue 479–480, 600–608. https://doi.org/10.1016/j.virol.2015.02.007

Ahmed, I., Buchert, R., Zhou, M., Jiao, X., Mittal, K., Sheikh, T.I., Scheller, U., Vasli, N., Rafiq, M.A., Brohi, M.Q., Mikhailov, A., Ayaz, M., Bhatti, A., Sticht, H., Nasr, T., Carter, M.T., Uebe, S., Reis, A., Ayub, M., John, P., Kiledjian, M., Vincent, J.B., Jamra, R.A., 2015. Mutations in DCPS and EDC3 in autosomal recessive intellectual disability indicate a crucial role for mRNA decapping in neurodevelopment. Hum. Mol. Genet. 24, 3172–3180. https://doi.org/10.1093/hmg/ddv069

Aizer, A., Kalo, A., Kafri, P., Shraga, A., Ben-Yishay, R., Jacob, A., Kinor, N., Shav-Tal, Y., 2014. Quantifying mRNA targeting to P-bodies in living human cells reveals their dual role in mRNA decay and storage. J. Cell Sci. 127, 4443–4456. https://doi.org/10.1242/jcs.152975

Alberti, S., Gladfelter, A., Mittag, T., 2019. Considerations and Challenges in Studying Liquid-Liquid Phase Separation and Biomolecular Condensates. Cell 176, 419–434. https://doi.org/10.1016/j.cell.2018.12.035

Arribas-Layton, M., Wu, D., Lykke-Andersen, J., Song, H., 2013. Structural and functional control of the eukaryotic mRNA decapping machinery. Biochim. Biophys. Acta BBA - Gene Regul. Mech., RNA Decay Mechanisms 1829, 580–589. https://doi.org/10.1016/j.bbagrm.2012.12.006

Badis, G., Saveanu, C., Fromont-Racine, M., Jacquier, A., 2004. Targeted mRNA Degradation by Deadenylation-Independent Decapping. Mol. Cell 15, 5–15. https://doi.org/10.1016/j.molcel.2004.06.028

Banani, S.F., Lee, H.O., Hyman, A.A., Rosen, M.K., 2017. Biomolecular condensates: organizers of cellular biochemistry. Nat. Rev. Mol. Cell Biol. 18, 285–298. https://doi.org/10.1038/nrm.2017.7

Banani, S.F., Rice, A.M., Peeples, W.B., Lin, Y., Jain, S., Parker, R., Rosen, M.K., 2016. Compositional Control of Phase-Separated Cellular Bodies. Cell 166, 651–663. https://doi.org/10.1016/j.cell.2016.06.010

Beelman, C.A., Stevens, A., Caponigro, G., LaGrandeur, T.E., Hatfield, L., Fortner, D.M., Parker, R., 1996. An essential component of the decapping enzyme required for normal rates of mRNA turnover. Nature 382, 642–646. https://doi.org/10.1038/382642a0

Boeynaems, S., Alberti, S., Fawzi, N.L., Mittag, T., Polymenidou, M., Rousseau, F., Schymkowitz, J., Shorter, J., Wolozin, B., Van Den Bosch, L., Tompa, P., Fuxreiter, M., 2018. Protein Phase Separation: A New Phase in Cell Biology. Trends Cell Biol. 28, 420–435. https://doi.org/10.1016/j.tcb.2018.02.004

Brengues, M., Teixeira, D., Parker, R., 2005. Movement of Eukaryotic mRNAs Between Polysomes and Cytoplasmic Processing Bodies. Science 310, 486–489. https://doi.org/10.1126/science.1115791

Chan, L.Y., Mugler, C.F., Heinrich, S., Vallotton, P., Weis, K., 2018. Non-invasive measurement of mRNA decay reveals translation initiation as the major determinant of mRNA stability. eLife 7, e32536. https://doi.org/10.7554/eLife.32536

Chang, Y.-F., Imam, J.S., Wilkinson, M.F., 2007. The Nonsense-Mediated Decay RNA Surveillance Pathway. Annu. Rev. Biochem. 76, 51–74. https://doi.org/10.1146/annurev.biochem.76.050106.093909

Charenton, C., Gaudon-Plesse, C., Fourati, Z., Taverniti, V., Back, R., Kolesnikova, O., Séraphin, B., Graille, M., 2017. A unique surface on Pat1 C-terminal domain directly interacts with Dcp2 decapping enzyme and Xrn1 5’–3’ mRNA exonuclease in yeast. Proc. Natl. Acad. Sci. 114, E9493–E9501. https://doi.org/10.1073/pnas.1711680114

Chen, I., Dorr, B.M., Liu, D.R., 2011. A general strategy for the evolution of bond-forming enzymes using yeast display. Proc. Natl. Acad. Sci. 108, 11399–11404. https://doi.org/10.1073/pnas.1101046108

Courchaine, E.M., Lu, A., Neugebauer, K.M., 2016. Droplet organelles? EMBO J. 35, 1603–1612. https://doi.org/10.15252/embj.201593517

Damman, R., Schütz, S., Luo, Y., Weingarth, M., Sprangers, R., Baldus, M., 2019. Atomic-level insight into mRNA processing bodies by combining solid and solution-state NMR spectroscopy. Nat. Commun. 10, 1–11. https://doi.org/10.1038/s41467-019-12402-3

Decker, C.J., Parker, R., 2012. P-Bodies and Stress Granules: Possible Roles in the Control of Translation and mRNA Degradation. Cold Spring Harb. Perspect. Biol. 4, a012286. https://doi.org/10.1101/cshperspect.a012286

Deshmukh, M.V., Jones, B.N., Quang-Dang, D.-U., Flinders, J., Floor, S.N., Kim, C., Jemielity, J., Kalek, M., Darzynkiewicz, E., Gross, J.D., 2008. mRNA Decapping Is Promoted by an RNA-Binding Channel in Dcp2. Mol. Cell 29, 324–336. https://doi.org/10.1016/j.molcel.2007.11.027

Dougherty, J.D., White, J.P., Lloyd, R.E., 2011. Poliovirus-Mediated Disruption of Cytoplasmic Processing Bodies. J. Virol. 85, 64–75. https://doi.org/10.1128/JVI.01657-10

Dunckley, T., Parker, R., 1999. The DCP2 protein is required for mRNA decapping in Saccharomyces cerevisiae and contains a functional MutT motif. EMBO J. 18, 5411–5422. https://doi.org/10.1093/emboj/18.19.5411

Franks, T.M., Lykke-Andersen, J., 2008. The Control of mRNA Decapping and P-Body Formation. Mol. Cell 32, 605–615. https://doi.org/10.1016/j.molcel.2008.11.001

Fromm, S.A., Kamenz, J., Nöldeke, E.R., Neu, A., Zocher, G., Sprangers, R., 2014. In Vitro Reconstitution of a Cellular Phase-Transition Process that Involves the mRNA Decapping Machinery. Angew. Chem. Int. Ed. 53, 7354–7359. https://doi.org/10.1002/anie.201402885

Fromm, S.A., Truffault, V., Kamenz, J., Braun, J.E., Hoffmann, N.A., Izaurralde, E., Sprangers, R., 2012. The structural basis of Edc3- and Scd6-mediated activation of the Dcp1:Dcp2 mRNA decapping complex. EMBO J. 31, 279–290. https://doi.org/10.1038/emboj.2011.408

Gallego, L.D., Schneider, M., Mittal, C., Romanauska, A., Carrillo, R.M.G., Schubert, T., Pugh, B.F., Köhler, A., 2020. Phase separation directs ubiquitination of gene-body nucleosomes. Nature 579, 592–597. https://doi.org/10.1038/s41586-020-2097-z

Harigaya, Y., Jones, B.N., Muhlrad, D., Gross, J.D., Parker, R., 2010. Identification and Analysis of the Interaction between Edc3 and Dcp2 in Saccharomyces cerevisiae. Mol. Cell. Biol. 30, 1446–1456. https://doi.org/10.1128/MCB.01305-09

He, F., Celik, A., Wu, C., Jacobson, A., 2018. General decapping activators target different subsets of inefficiently translated mRNAs. eLife 7, e34409. https://doi.org/10.7554/eLife.34409

He, F., Jacobson, A., 2015. Control of mRNA decapping by positive and negative regulatory elements in the Dcp2 C-terminal domain. RNA 21, 1633–1647. https://doi.org/10.1261/rna.052449.115

He, F., Li, C., Roy, B., Jacobson, A., 2014. Yeast Edc3 Targets RPS28B mRNA for Decapping by Binding to a 3’ Untranslated Region Decay-Inducing Regulatory Element. Mol. Cell. Biol. 34, 1438–1451. https://doi.org/10.1128/MCB.01584-13

Hnisz, D., Shrinivas, K., Young, R.A., Chakraborty, A.K., Sharp, P.A., 2017. A Phase Separation Model for Transcriptional Control. Cell 169, 13–23. https://doi.org/10.1016/j.cell.2017.02.007

Hondele, M., Sachdev, R., Heinrich, S., Wang, J., Vallotton, P., Fontoura, B.M.A., Weis, K., 2019. DEAD-box ATPases are global regulators of phase-separated organelles. Nature 573, 144–148. https://doi.org/10.1038/s41586-019-1502-y

Horvathova, I., Voigt, F., Kotrys, A.V., Zhan, Y., Artus-Revel, C.G., Eglinger, J., Stadler, M.B., Giorgetti, L., Chao, J.A., 2017. The Dynamics of mRNA Turnover Revealed by Single-Molecule Imaging in Single Cells. Mol. Cell 68, 615–625.e9. https://doi.org/10.1016/j.molcel.2017.09.030

Hubstenberger, A., Courel, M., Bénard, M., Souquere, S., Ernoult-Lange, M., Chouaib, R., Yi, Z., Morlot, J.-B., Munier, A., Fradet, M., Daunesse, M., Bertrand, E., Pierron, G., Mozziconacci, J., Kress, M., Weil, D., 2017. P-Body Purification Reveals the Condensation of Repressed mRNA Regulons. Mol. Cell 68, 144–157.e5. https://doi.org/10.1016/j.molcel.2017.09.003

Hyman, A.A., Weber, C.A., Jülicher, F., 2014. Liquid-Liquid Phase Separation in Biology. Annu. Rev. Cell Dev. Biol. 30, 39–58. https://doi.org/10.1146/annurev-cellbio-100913-013325

Jonas, S., Izaurralde, E., 2013. The role of disordered protein regions in the assembly of decapping complexes and RNP granules. Genes Dev. 27, 2628–2641. https://doi.org/10.1101/gad.227843.113

Keenen, M.M., Larson, A.G., Narlikar, G.J., 2018. Chapter Three - Visualization and Quantitation of Phase-Separated Droplet Formation by Human HP1α, in: Rhoades, E. (Ed.), Methods in Enzymology, Intrinsically Disordered Proteins. Academic Press, pp. 51–66. https://doi.org/10.1016/bs.mie.2018.09.034

Kim, T.H., Tsang, B., Vernon, R.M., Sonenberg, N., Kay, L.E., Forman-Kay, J.D., 2019. Phospho-dependent phase separation of FMRP and CAPRIN1 recapitulates regulation of translation and deadenylation. Science 365, 825–829. https://doi.org/10.1126/science.aax4240

Kowalska, J., Osowniak, A., Zuberek, J., Jemielity, J., 2012. Synthesis of nucleoside phosphosulfates. Bioorg. Med. Chem. Lett. 22, 3661–3664. https://doi.org/10.1016/j.bmcl.2012.04.039

Lee, W., Tonelli, M., Markley, J.L., 2015. NMRFAM-SPARKY: enhanced software for biomolecular NMR spectroscopy. Bioinformatics 31, 1325–1327. https://doi.org/10.1093/bioinformatics/btu830

Li, Y., Dai, J., Song, M., Fitzgerald-Bocarsly, P., Kiledjian, M., 2012. Dcp2 Decapping Protein Modulates mRNA Stability of the Critical Interferon Regulatory Factor (IRF) IRF-7. Mol. Cell. Biol. 32, 1164–1172. https://doi.org/10.1128/MCB.06328-11

Lobel, J.H., Tibble, R.W., Gross, J.D., 2019. Pat1 activates late steps in mRNA decay by multiple mechanisms. Proc. Natl. Acad. Sci. 116, 23512–23517. https://doi.org/10.1073/pnas.1905455116

Moore, M.J., 2005. From Birth to Death: The Complex Lives of Eukaryotic mRNAs. Science 309, 1514–1518. https://doi.org/10.1126/science.1111443

Mugler, C.F., Hondele, M., Heinrich, S., Sachdev, R., Vallotton, P., Koek, A.Y., Chan, L.Y., Weis, K., 2016. ATPase activity of the DEAD-box protein Dhh1 controls processing body formation. eLife 5, e18746. https://doi.org/10.7554/eLife.18746

Mugridge, J.S., Coller, J., Gross, J.D., 2018. Structural and molecular mechanisms for the control of eukaryotic 5’–3’ mRNA decay. Nat. Struct. Mol. Biol. 25, 1077–1085. https://doi.org/10.1038/s41594-018-0164-z

Müller-McNicoll, M., Neugebauer, K.M., 2013. How cells get the message: dynamic assembly and function of mRNA–protein complexes. Nat. Rev. Genet. 14, 275–287. https://doi.org/10.1038/nrg3434

Nagarajan, V.K., Jones, C.I., Newbury, S.F., Green, P.J., 2013. XRN 5’→3’ exoribonucleases: Structure, mechanisms and functions. Biochim. Biophys. Acta BBA - Gene Regul. Mech., RNA Decay Mechanisms 1829, 590–603. https://doi.org/10.1016/j.bbagrm.2013.03.005

Paquette, D.R., Tibble, R.W., Daifuku, T.S., Gross, J.D., 2018. Control of mRNA decapping by autoinhibition. Nucleic Acids Res. 46, 6318–6329. https://doi.org/10.1093/nar/gky233

Parker, R., Sheth, U., 2007. P Bodies and the Control of mRNA Translation and Degradation. Mol. Cell 25, 635–646. https://doi.org/10.1016/j.molcel.2007.02.011

Pitchiaya, S., Mourao, M.D.A., Jalihal, A.P., Xiao, L., Jiang, X., Chinnaiyan, A.M., Schnell, S., Walter, N.G., 2019. Dynamic Recruitment of Single RNAs to Processing Bodies Depends on RNA Functionality. Mol. Cell 74, 521–533.e6. https://doi.org/10.1016/j.molcel.2019.03.001

Rao, B.S., Parker, R., 2017. Numerous interactions act redundantly to assemble a tunable size of P bodies in Saccharomyces cerevisiae. Proc. Natl. Acad. Sci. 114, E9569–E9578. https://doi.org/10.1073/pnas.1712396114

Refaei, M.A., Combs, A., Kojetin, D.J., Cavanagh, J., Caperelli, C., Rance, M., Sapitro, J., Tsang, P., 2011. Observing selected domains in multi-domain proteins via sortase-mediated ligation and NMR spectroscopy. J. Biomol. NMR 49, 3–7. https://doi.org/10.1007/s10858-010-9464-2

Riback, J.A., Zhu, L., Ferrolino, M.C., Tolbert, M., Mitrea, D.M., Sanders, D.W., Wei, M.-T., Kriwacki, R.W., Brangwynne, C.P., 2020. Composition-dependent thermodynamics of intracellular phase separation. Nature 1–6. https://doi.org/10.1038/s41586-020-2256-2

Sanulli, S., Trnka, M.J., Dharmarajan, V., Tibble, R.W., Pascal, B.D., Burlingame, A.L., Griffin, P.R., Gross, J.D., Narlikar, G.J., 2019. HP1 reshapes nucleosome core to promote phase separation of heterochromatin. Nature 575, 390–394. https://doi.org/10.1038/s41586-019-1669-2

Schindelin, J., Arganda-Carreras, I., Frise, E., Kaynig, V., Longair, M., Pietzsch, T., Preibisch, S., Rueden, C., Saalfeld, S., Schmid, B., Tinevez, J.-Y., White, D.J., Hartenstein, V., Eliceiri, K., Tomancak, P., Cardona, A., 2012. Fiji: an open-source platform for biological-image analysis. Nat. Methods 9, 676–682. https://doi.org/10.1038/nmeth.2019

Schütz, S., Nöldeke, E.R., Sprangers, R., 2017. A synergistic network of interactions promotes the formation of in vitro processing bodies and protects mRNA against decapping. Nucleic Acids Res. 45, 6911–6922. https://doi.org/10.1093/nar/gkx353

She, M., Decker, C.J., Svergun, D.I., Round, A., Chen, N., Muhlrad, D., Parker, R., Song, H., 2008. Structural Basis of Dcp2 Recognition and Activation by Dcp1. Mol. Cell 29, 337–349. https://doi.org/10.1016/j.molcel.2008.01.002

Sheth, U., Parker, R., 2003. Decapping and Decay of Messenger RNA Occur in Cytoplasmic Processing Bodies. Science 300, 805–808. https://doi.org/10.1126/science.1082320

Sheu-Gruttadauria, J., MacRae, I.J., 2018. Phase Transitions in the Assembly and Function of Human miRISC. Cell 173, 946–957.e16. https://doi.org/10.1016/j.cell.2018.02.051

Shukla, S., Parker, R., 2014. Quality control of assembly-defective U1 snRNAs by decapping and 5’-to-3’ exonucleolytic digestion. Proc. Natl. Acad. Sci. 111, E3277–E3286. https://doi.org/10.1073/pnas.1412614111

Teixeira, D., Parker, R., 2007. Analysis of P-Body Assembly in Saccharomyces cerevisiae. Mol. Biol. Cell 18, 2274–2287. https://doi.org/10.1091/mbc.e07-03-0199

Trotman, J.B., Schoenberg, D.R., 2019. A recap of RNA recapping. WIREs RNA 10, e1504. https://doi.org/10.1002/wrna.1504

Tutucci, E., Vera, M., Biswas, J., Garcia, J., Parker, R., Singer, R.H., 2018. An improved MS2 system for accurate reporting of the mRNA life cycle. Nat. Methods 15, 81–89. https://doi.org/10.1038/nmeth.4502

Van Treeck, B., Parker, R., 2018. Emerging Roles for Intermolecular RNA-RNA Interactions in RNP Assemblies. Cell 174, 791–802. https://doi.org/10.1016/j.cell.2018.07.023

Wang, C., Schmich, F., Srivatsa, S., Weidner, J., Beerenwinkel, N., Spang, A., 2018. Context-dependent deposition and regulation of mRNAs in P-bodies. eLife 7, e29815. https://doi.org/10.7554/eLife.29815

Wang, Z., Jiao, X., Carr-Schmid, A., Kiledjian, M., 2002. The hDcp2 protein is a mammalian mRNA decapping enzyme. Proc. Natl. Acad. Sci. 99, 12663–12668. https://doi.org/10.1073/pnas.192445599

Warminski, M., Sikorski, P.J., Warminska, Z., Lukaszewicz, M., Kropiwnicka, A., Zuberek, J., Darzynkiewicz, E., Kowalska, J., Jemielity, J., 2017. Amino-Functionalized 5’ Cap Analogs as Tools for Site-Specific Sequence-Independent Labeling of mRNA. Bioconjug. Chem. 28, 1978–1992. https://doi.org/10.1021/acs.bioconjchem.7b00291

Wurm, J.P., Holdermann, I., Overbeck, J.H., Mayer, P.H.O., Sprangers, R., 2017. Changes in conformational equilibria regulate the activity of the Dcp2 decapping enzyme. Proc. Natl. Acad. Sci. 114, 6034–6039. https://doi.org/10.1073/pnas.1704496114

Wurm, J.P., Overbeck, J., Sprangers, R., 2016. The *S. pombe mRNA decapping complex recruits cofactors and an Edc1-like activator through a single dynamic surface*. RNA 22, 1360–1372. https://doi.org/10.1261/rna.057315.116

Xing, W., Muhlrad, D., Parker, R., Rosen, M.K., 2018. A quantitative inventory of yeast P body proteins reveals principles of compositional specificity. bioRxiv 489658. https://doi.org/10.1101/489658

