## Supplemental Materials for "Biomolecular condensates amplify mRNA decapping by coupling protein interactions with conformational changes in Dcp1/Dcp2"

#### Supplementary Materials

**Supplemental Figures** **Page**

**Figure S1:** Droplet behavior of Dcp1/$Dcp2_{ext}$ and Dcp1/$Dcp2_{ext}$/Edc3 (Related to Figure 1) **5**
**Figure S2:** Synthesis of dual-labeled 35mer RNA and probe stability over course of decapping assay (Related to Figure 3) **6**
**Figure S3:** Monitoring decapping of dual-labeled 35mer RNA using fluorescence polarization (Related to Figure 3) **7**
**Figure S4:** Activation of Dcp1/$Dcp2_{ext}$ by Edc3(Lsm) and Dcp1/$Dcp2_{ext}$ depletion assay (Related to Figure 4) **8**
**Figure S5:** Enrichment of Dcp1/$Dcp2_{ext}$ and RNA in liquid droplets at varying Edc3 conditions (Related to Figure 4) **9**
**Figure S6:** SortaseA-mediated ligation of Dcp2 (Related to Figure 5) **10**
**Figure S7:** $^{1}H/^{13}C$-HSQC of ligated Dcp1/$Dcp2_{ext}$ in absence and presence of Edc3 (Related to Figure 5) **11**
**Figure S8:** Representative resonances reporting on fast equilibrium between open and inactive states (Related to Figure 5A, 5B & 5C) **12**
**Figure S9:** Comparison of NMR chemical shift perturbations of Dcp1/$Dcp2_{ext}$ and Dcp1/$Dcp2_{ext}$/Edc3 relative to Dcp1/$Dcp2_{core}$ (Related to Figure 5B) **13**
**Figure S10:** Fluorescence polarization curves and fit parameters from Dcp1/$Dcp2_{ext}$ and Dcp1/$Dcp2_{ext}$(Y220G) decapping assays (Related to Figure 5C & 5D) **14**

**Supplementary Data Tables**

**Table 1:** Protein constructs used in this study **15**
**Table 2:** Relative activity of Dcp1/$Dcp2_{ext}$ in liquid droplets with or without Edc3 **16**
**Table 3:** Relative activity of Dcp1/$Dcp2_{ext}$ in Bulk and Supernatant with variable Edc3 concentrations **16**
**Table 4:** Relative activity of Dcp1/$Dcp2_{ext}$ upon addition of varying Edc3(Lsm) domain **16**
**Table 5:** Relative activity of Dcp1/$Dcp2_{ext}$(Y220G) in Bulk and Supernatant with variable Edc3 concentrations **16**
**Table 6:** Equilibrium dissociation constants ($K_D$) for various Dcp2 constructs determined from fluorescence polarization **17**

**Figure S1**. Dcp1/Dcp2$_{ext}$/Edc3 droplets are more homogenous in size and do not always undergo homogenous mixing. **A**. Fusion of Dcp1/Dcp2$_{ext}$ droplets. Each image is taken approximately one minute after the preceding. **B**. Fusion of Dcp1/Dcp2$_{ext}$/Edc3 droplets. Each image is taken approximately one minute after the preceding image. **C**. Brightfield images of serial two-fold diluted Dcp2$_{ext}$ show it forms small liquid droplets at high concentrations. **D**. Brightfield images of the disordered C-terminus show it is sufficient to undergo phase separation in a concentration-dependent manner. **E**. (Left) Histogram of droplet area (in $\mu m^2$) observed for Dcp1/Dcp2$_{ext}$ and stoichiometric Dcp1/Dcp2$_{ext}$/Edc3 droplets at concentrations 10-fold greater than $c_{sat}$, the concentration at which phase separation occurs. (Right) Cumulative Distribution Function of the data presented on the left demonstrates that Dcp1/Dcp2$_{ext}$ droplets exhibit a much wider distribution of sizes and cannot be adequately explained by only droplets ≤ 100 $\mu m^2$ . Scale bar in **A**, **B**, 10 $\mu$m and in **C**, **D**, 20 $\mu$m.

**Figure S2**. Synthesis and stability of dually labelled 5' capped 35 nt RNA probe. A) Overview of the labelling procedure, IVT – in vitro transcription, c.t. – co-transcriptional, p.t. – post-transcriptional; B) structures of the reagents used for the 5' and 3' end labelling; C) Analysis of purified 5' capped & labelled RNA after IVT: lane 1 – reference uncapped RNA, lane 2 – RNA capped co-transcriptionally with fluorescent cap analog (FAM-m$^7$Gp$_3$A$_m$pG). Conditions of PAGE analysis: 15% polyacrylamide, 7M urea in TBE, 40 ng RNA, the gel was visualized by fluorescein emission (FAM) followed by staining with Ethidium Bromide (EtBr); D) Labelling of the 3' end of FAM-m$^7$Gp$_3$A$_m$pG-RNA with pAp-SCy5 to yield dually labelled probe: lane 3 – crude dually labelled RNA after purification with the Clean and Concentrator-5 kit; lane 4 – HPLC-purified RNA probe. Conditions of PAGE analysis are as above, with exception that additional visualization by Cy5 emission was performed. **E**. Localization of both dyes of dually labelled RNA probe to Dcp1/Dcp2$_{ext}$/Edc3 droplets in the absence of $MgCl_2$ over twenty minutes. **F**. Quantification of intensity for both the m7G cap (fluorescein) and RNA body (Cy5) at time points in E shows no significant loss in either fluorescein or Cy5 intensity over time.

**Figure S3**. Monitoring decapping of dual-labeled 35mer RNA using fluorescence polarization. **A.** Kinetic trace demonstrating release of FAM-m$^7$GDP as a result of decapping by Dcp1/Dcp2$_{core}$, which leads to a time-dependent decrease in mP from more rapid tumbling of released FAM-m$^7$GDP. **B**. Observed rates of decapping determined from **A** demonstrate Dcp1/Dcp2$_{core}$ is able to hydrolyze dual-labeled substrate at 0.2 min$^{-1}$.

**Figure S4**. Edc3 sequesters Dcp1/Dcp2$_{ext}$ in condensates to cooperatively activate decapping. **A**. Concentration-dependent enhancement of decapping by the Lsm domain of Edc3 occurs in the absence of microscopically visible phase-separated condensates at any concentration of the Lsm domain (bottom panel of brightfield images). **B**. Hill coefficients from best-fit of curves for Dcp1/Dcp2$_{ext}$/Edc3 from Figure 3C and Dcp1/Dcp2$_{ext}$/Edc3(Lsm domain) in S3A.

Dimeric Edc3 results in a five-fold increase in cooperativity of activation of decapping relative to the Lsm domain. **C**. Depletion of Dcp1/Dcp2$_{ext}$ from the supernatant at concentrations of Edc3 ranging from 78.1 nM to 80 μM visualized by SDS-PAGE gel. (Top) Instant Blue staining reveals total protein levels in the supernatant following pelleting of liquid droplets (see Methods). (Bottom) In-gel fluorescence of Cy5-labelled Dcp2 shows its Edc3-dependent disappearance from the solution. **D.** Quantification of Dcp1/Dcp2$_{ext}$ from analysis of Dcp2 stained with Instant Blue (dark red) or measured by in-gel Cy5 emission (pink triangles). Cy5 fluorescence is reproduced from main text Figure 3D for comparative purposes.

**Figure S5**. Composition of Dcp1/Dcp2$_{ext}$/Edc3 droplets alters Dcp1/Dcp2$_{ext}$ enrichment in phases. **A**. Enrichment of Dcp1/Dcp2$_{ext}$ in droplets of varying composition of Dcp1/Dcp2$_{ext}$ and Edc3. Dcp1/Dcp2$_{ext}$ enrichment is relatively constant in droplets in the absence of RNA (orange bars) and is moderately affected by the addition of RNA (blue bars). The addition of excess Edc3, however, greatly increases the enrichment of Dcp1/Dcp2$_{ext}$ in droplets and the addition of RNA further enriches the decapping complex in droplets. Images below graph correspond to Cy5-labelled Dcp1/Dcp2$_{ext}$. Error bars represent standard error of the mean. **B**. Enrichment of FAM-29mer RNA in droplets of varying Dcp1/Dcp2$_{ext}$ and Edc3 ratios. Error bars are s.e.m. Scale bar, 10 μm.

**Figure S6**. Generation of segmentally-labeled Dcp1/Dcp2$_{ext}$ for NMR. **A.** Reaction scheme for ligation of ILVMA-labeled, NMR active Dcp1/Dcp2 core domains with NMR invisible C-terminus using an enhanced SortaseA (eSrtA, See Methods). **B**. SDS-PAGE for the ligation reaction. The left gel corresponds to the reaction and purification of ligation product while wild-type Dcp1/Dcp2$_{ext}$ is shown on the right gel image. *Indicates N-terminal MBP tag generated upon incubation with TEV. **Indicates unreacted C-terminus generated upon incubation with TEV. ***Indicates eSrtA band. **C**. Ligation product and wild-type Dcp1/Dcp2$_{ext}$ exhibit similar decapping activity on a capped 29mer substrate.

**Figure S7**. 1H/13C-HSQC of ligated Dcp1/Dcp2$_{ext}$. **A**. Dcp1/Dcp2 core domains (orange), Dcp1/Dcp2$_{ext}$ (red), and Dcp1/Dcp2$_{ext}$/Edc3 (green) spectra are overlaid. Assignments correspond to positions for core domains of Dcp1/Dcp2 and demonstrate that numerous assignments can be reliably transferred to Dcp1/Dcp2$_{ex}$. It should be noted several resonances broaden or undergo significant perturbations upon ligation of the C-terminus, suggesting significant structural rearrangements occur.

**Figure S8**. Several resonances in the catalytic domain of Dcp2 report on the open-to-inactive equilibrium. **A**. Additional $^{1}H/^{13}C$-methyl resonances used to calculate the population of the inactive state resulting from the addition of regulatory elements. Chemical shifts for the various constructs fall along a linear trajectory (dotted line), indicative of fast interconversion between the two states on the NMR timescale.

**Figure S9**. Addition of Edc3 reduces chemical shift perturbations (CSPs) of Dcp1/Dcp2$_{ext}$ to more closely resemble the chemical shifts of Dcp1/Dcp2$_{core}$. **A**. CSPs of Dcp1/Dcp2$_{ext}$ containing the C-terminus in the absence (red) and presence (green) of Edc3 for ILVMA terminal methyl resonances in the structured catalytic core of Dcp2. CSPs are calculated relative to Dcp1/Dcp2$_{core}$, which lacks the disordered C-terminus. The strong reduction in CSPs upon the addition of Edc3 indicates the structure of the core domains more closely resembles that of the uninhibited Dcp1/Dcp2 complex. The dotted line represents the standard deviation of observed CSPs plotted in the graph. **B**. (Left) CSPs for Dcp1/Dcp2$_{ext}$ mapped onto the inactive state of Dcp1/Dcp2 (PDB: 2qkm). (Right) CSPs for Dcp1/Dcp2$_{ext}$/Edc3 mapped onto the inactive state. ILVMA residues are shown as spheres. Dcp1 is shown as the yellow ribbon structure.

**Figure S10***.* The ability of Dcp2 to interact with RNA can be tuned across several orders of magnitude and the Y220G mutation in Dcp2 activates substrate binding and reduces the ability of Edc3 to stimulate decapping. **A**. Normalized FP curves for various Dcp2 constructs binding to U30mer RNA. **B**. The Y220G mutation does not affect the cooperativity of activation by Edc3 but increases the $K_{1/2}$ of activation three-fold. Error bars represent s.e.m. in **A** and standard error of the fit in **B**.

#### Figure S1

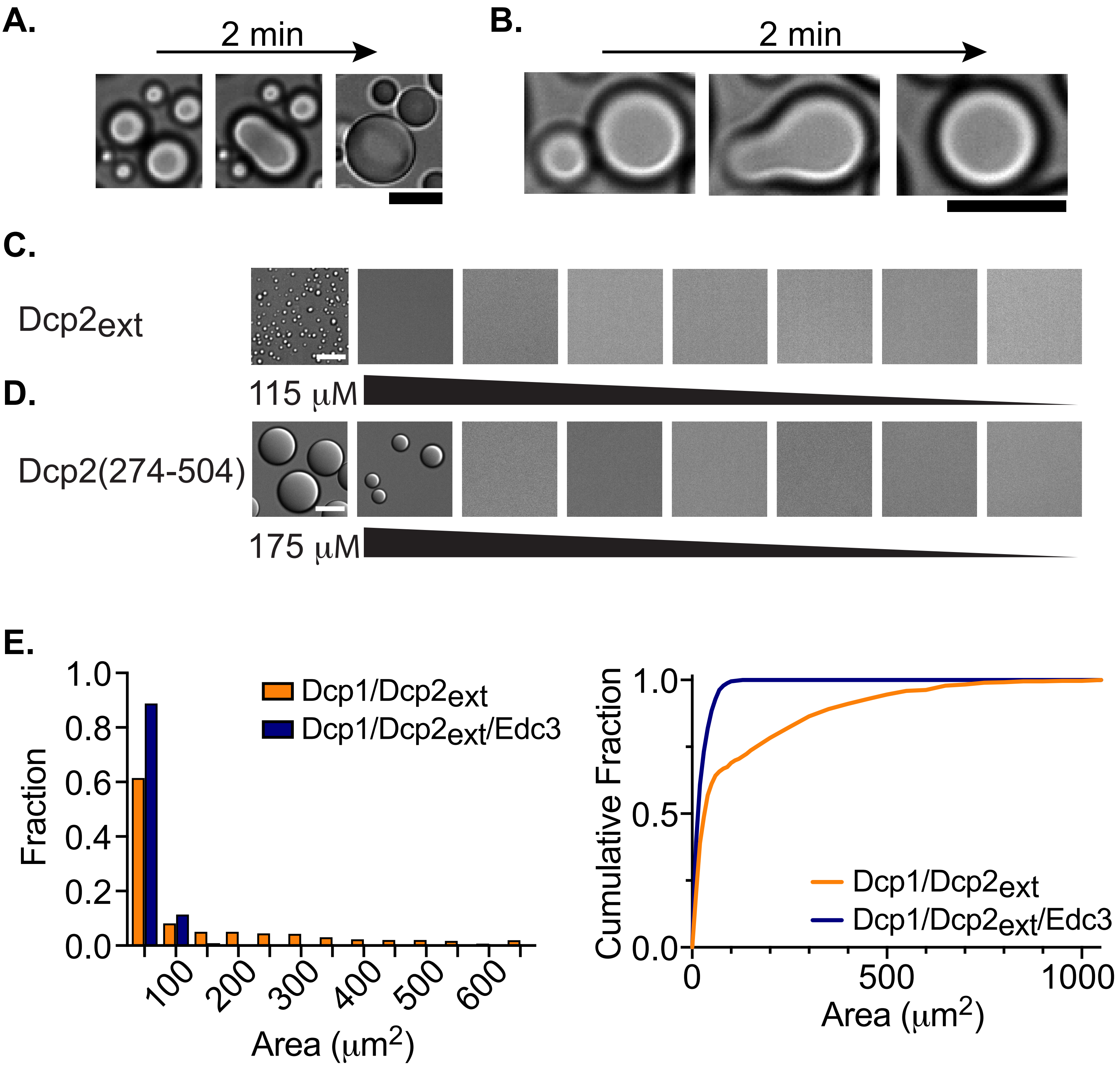

A.
2 min
B.
2 min
C.
Dcp2ext
D.
115 μM
Dcp2(274-504)
175 μM
E.
1.0
0.8
0.6
0.4
0.2
0.0
Fraction
Dcp1/Dcp2ext
Dcp1/Dcp2ext/Edc3
100
200
300
400
500
600
Area (μm2)
Cumulative Fraction
1.0
0.5
0.0
0
500
1000
Area (μm2)
Dcp1/Dcp2ext
Dcp1/Dcp2ext/Edc3

#### Figure S2

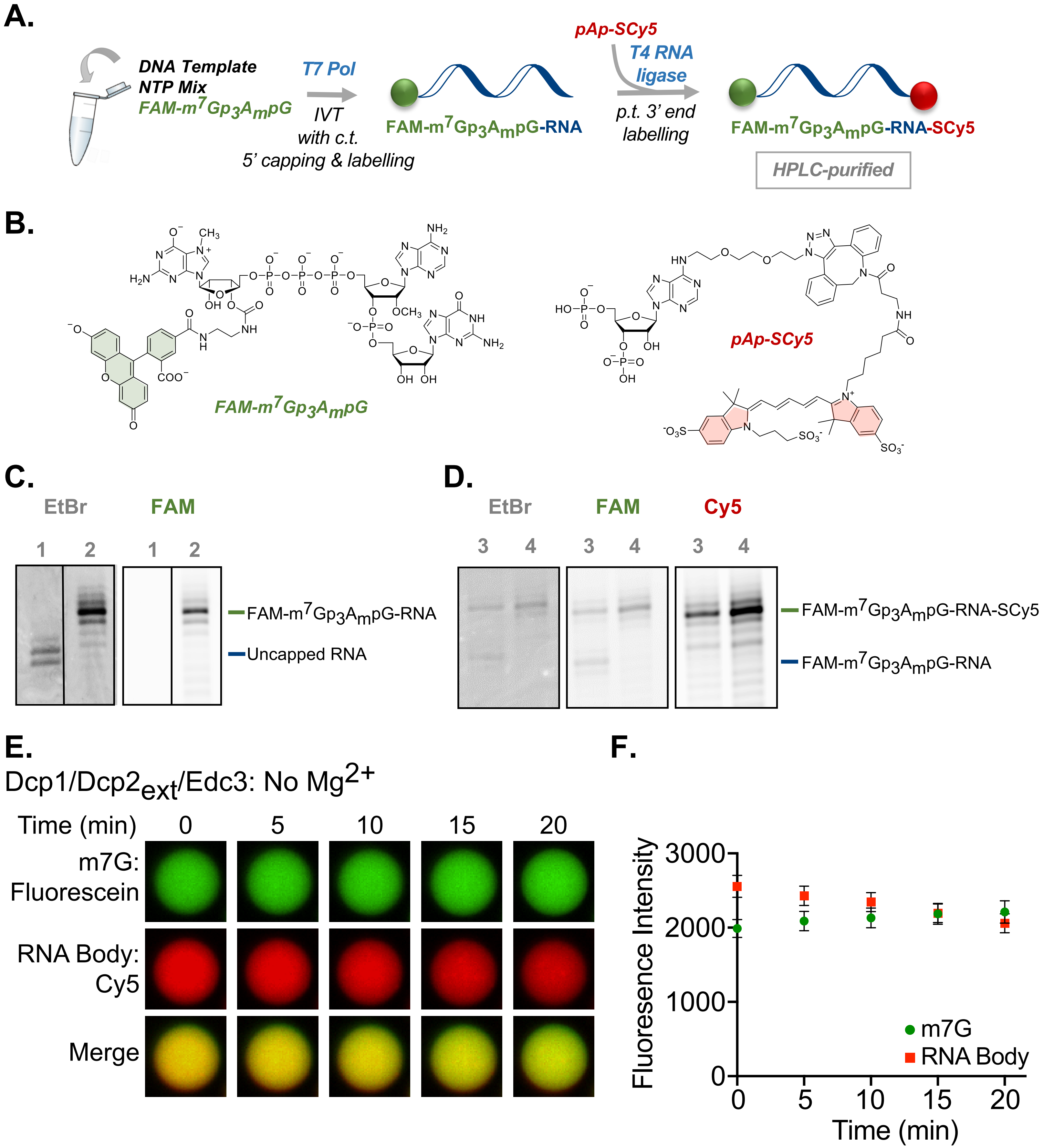
A.
DNA Template
NTP Mix
FAM-m7Gp3AmpG
T7 Pol
IVT
with c.t.
5' capping & labelling
FAM-m7Gp3AmpG-RNA
pAp-SCy5
T4 RNA ligase
p.t. 3' end labelling
FAM-m7Gp3AmpG-RNA-SCy5
HPLC-purified
B.
FAM-m7Gp3AmpG
pAp-SCy5
C.
EtBr
FAM
1 2 1 2
FAM-m7Gp3AmpG-RNA
Uncapped RNA
D.
EtBr
FAM
Cy5
3 4 3 4 3 4
FAM-m7Gp3AmpG-RNA-SCy5
FAM-m7Gp3AmpG-RNA
E.
Dcp1/Dcp2ext/Edc3: No Mg2+
Time (min) 0 5 10 15 20
m7G: Fluorescein
RNA Body: Cy5
Merge
F.
Fluorescence Intensity
3000
2000
1000
0
0 5 10 15 20
Time (min)
m7G
RNA Body

#### Figure S3

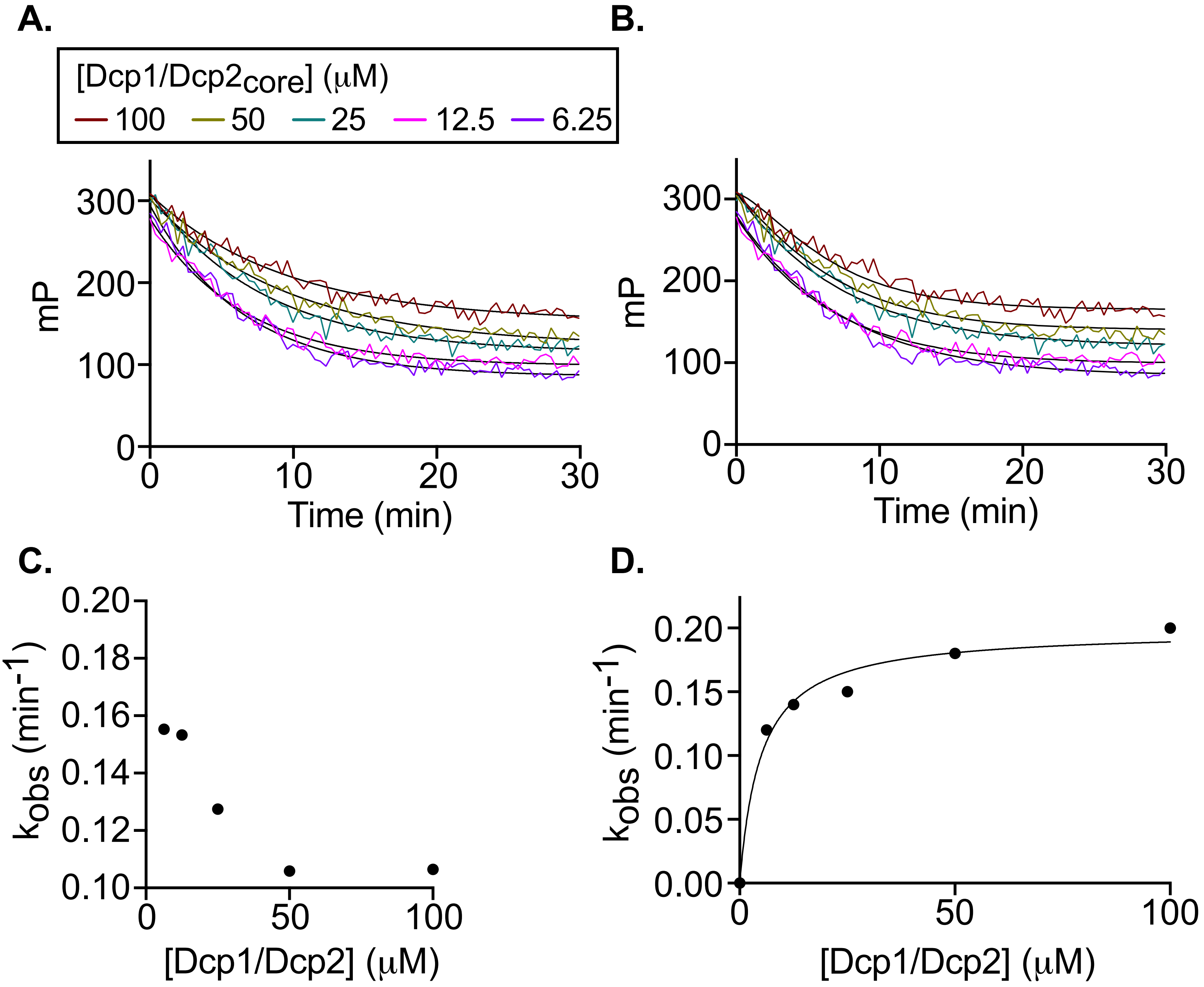
A.
B.
C.
D.
[Dcp1/Dcp2core] (μM)
100
50
25
12.5
6.25
mP
Time (min)
0
100
200
300
10
20
30
kobs (min-1)
0.10
0.12
0.14
0.16
0.18
0.20
0.00
0.05
0.15
[Dcp1/Dcp2] (μM)
50

**Figure S4**

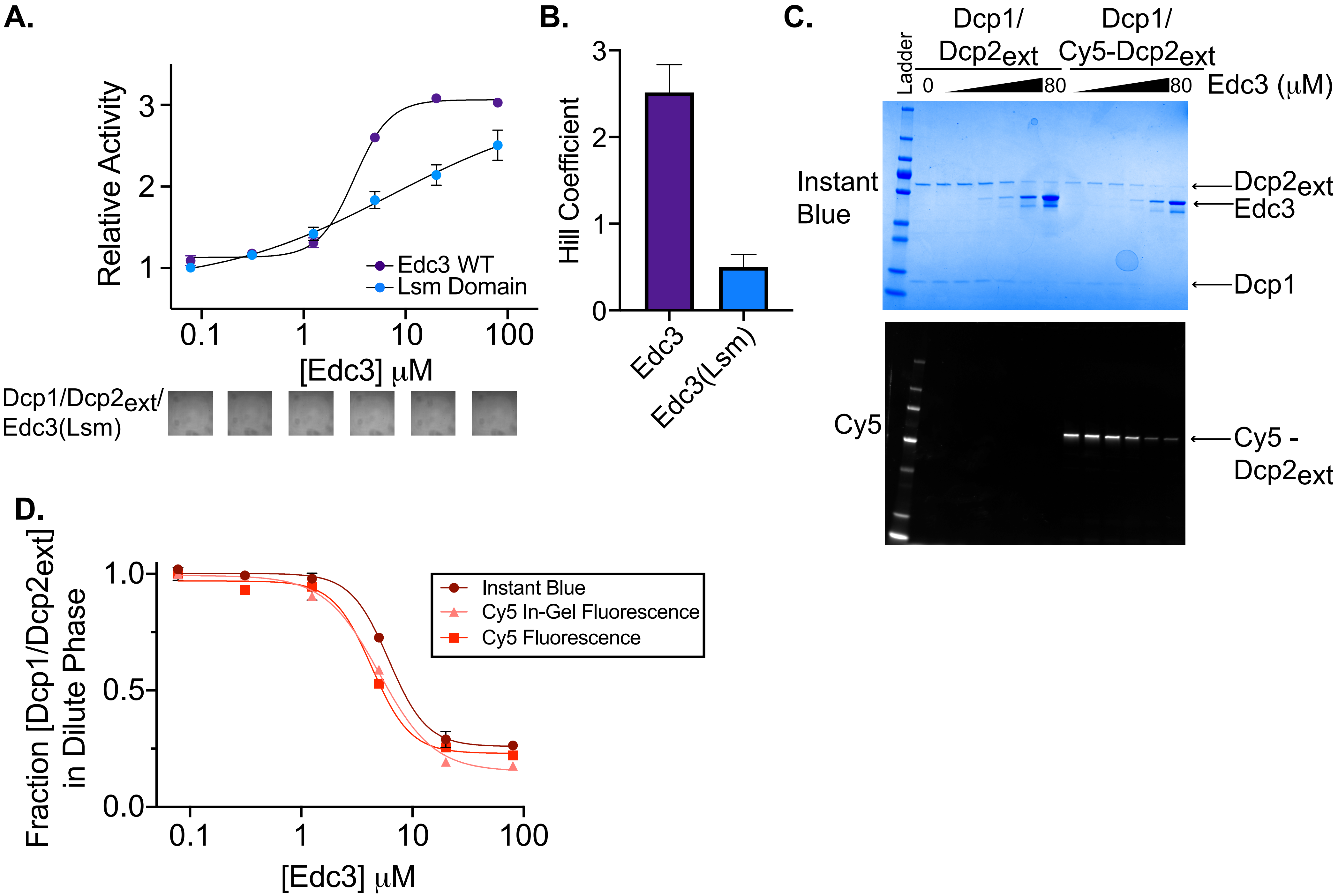
A.
Relative Activity
[Edc3] μM
Edc3 WT
Lsm Domain
Dcp1/Dcp2ext/
Edc3(Lsm)
B.
Hill Coefficient
Edc3
Edc3(Lsm)
C.
Ladder
Dcp1/
Dcp2ext
Dcp1/
Cy5-Dcp2ext
0
80
80
Edc3 (μM)
Instant
Blue
Dcp2ext
Edc3
Dcp1
Cy5
Cy5 -
Dcp2ext
D.
Fraction [Dcp1/Dcp2ext]
in Dilute Phase
[Edc3] μM
Instant Blue
Cy5 In-Gel Fluorescence
Cy5 Fluorescence

#### Figure S5

**A.**

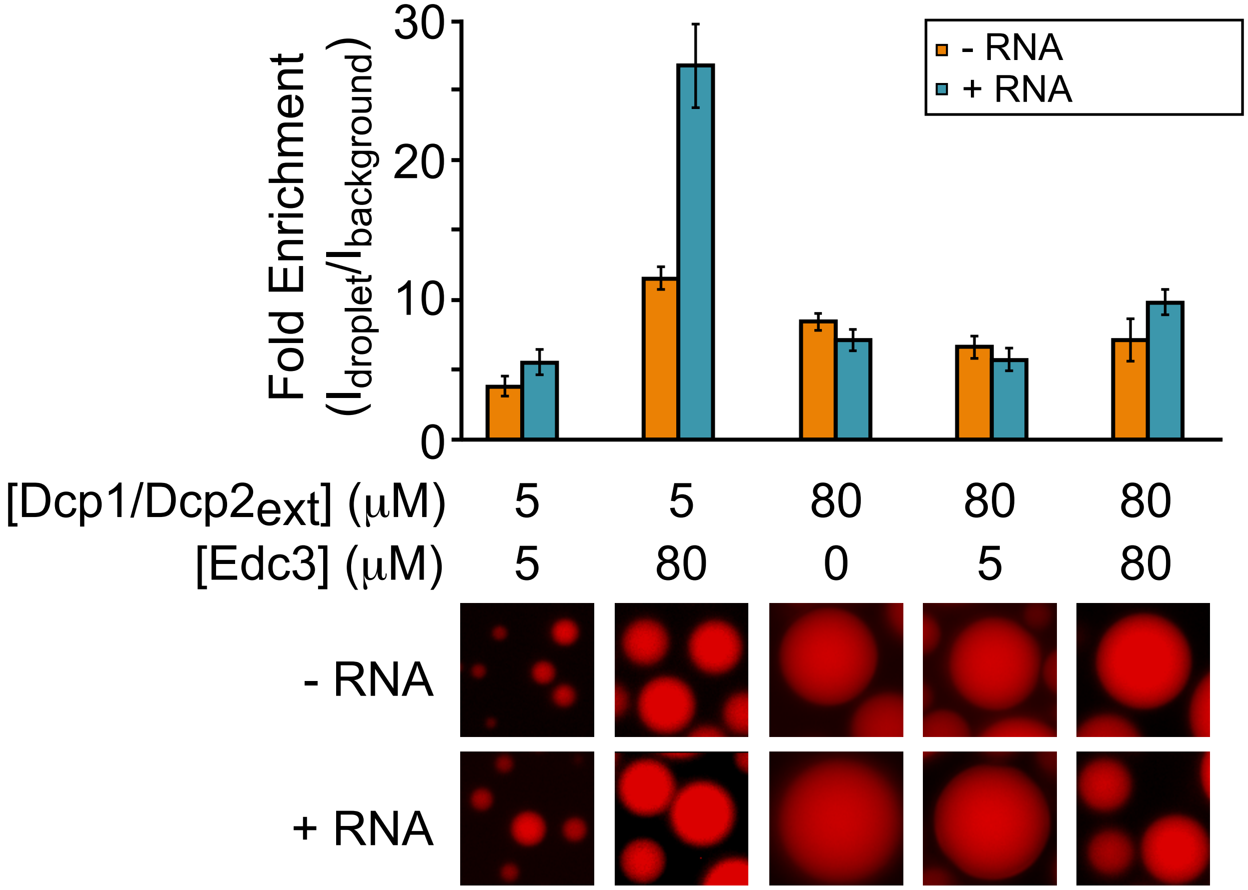

Fold Enrichment
($I_{droplet}/I_{background}$)
30
20
10
0
- RNA
+ RNA
[Dcp1/Dcp2ext] (μM) 5 5 80 80 80
[Edc3] (μM) 5 80 0 5 80
- RNA
+ RNA


**B.**

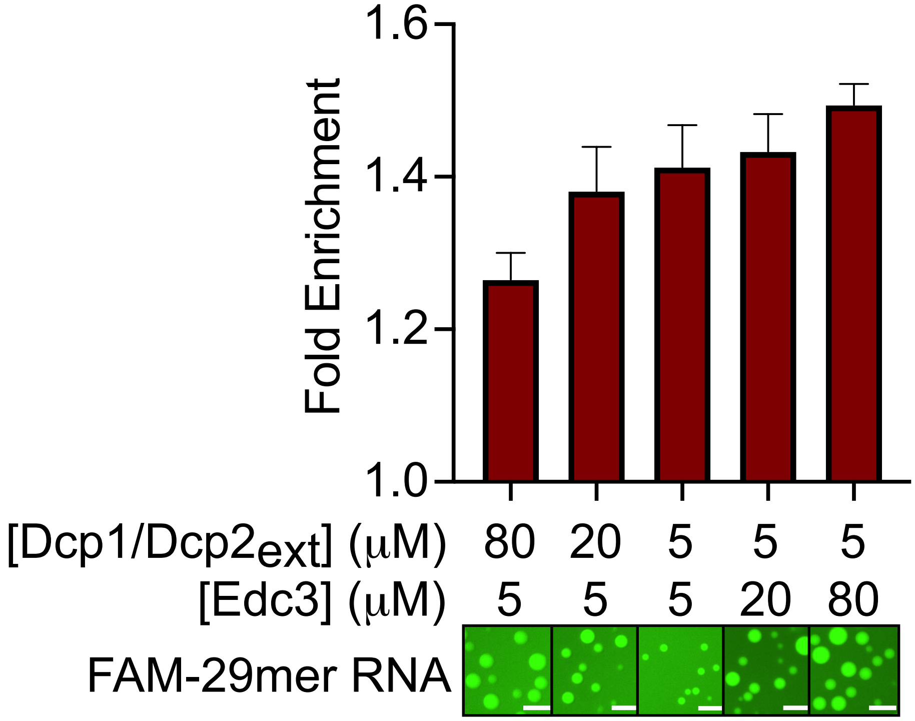

Fold Enrichment
1.6
1.4
1.2
1.0
[Dcp1/Dcp2ext] (μM) 80 20 5 5 5
[Edc3] (μM) 5 5 5 20 80
FAM-29mer RNA

### Figure S6

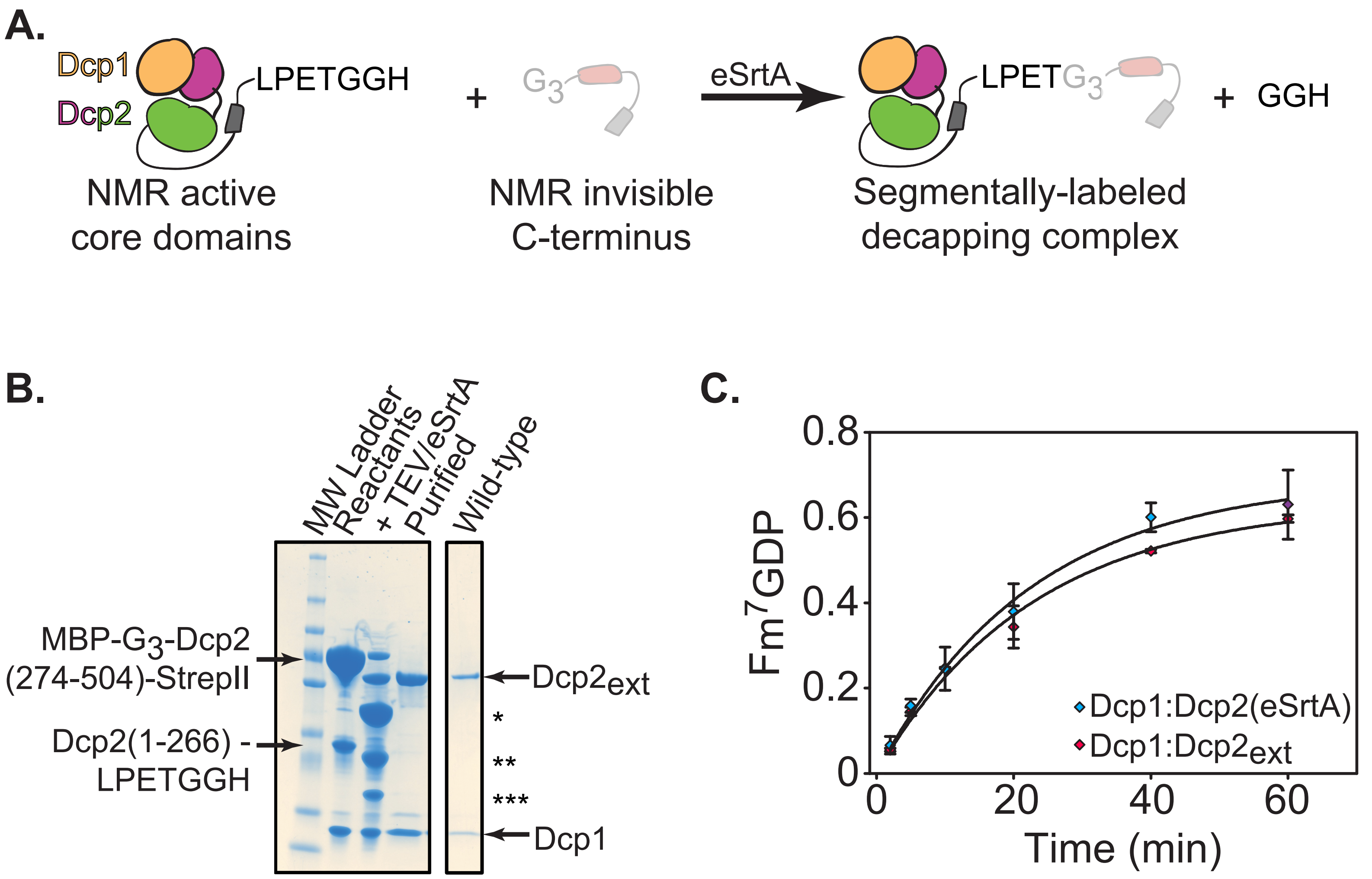
A.
Dcp1
Dcp2
LPETGGH
NMR active
core domains
+
G3
NMR invisible
C-terminus
eSrtA
LPETG3
Segmentally-labeled
decapping complex
+ GGH
B.
MW Ladder
Reactants
+ TEV/eSrtA
Purified
Wild-type
MBP-G3-Dcp2
(274-504)-StrepII
Dcp2(1-266) -
LPETGGH
Dcp2ext
*
**
***
Dcp1
C.
Fm7GDP
0.8
0.6
0.4
0.2
0
0
20
40
60
Time (min)
Dcp1:Dcp2(eSrtA)
Dcp1:Dcp2ext

**Figure S7**

**A.**

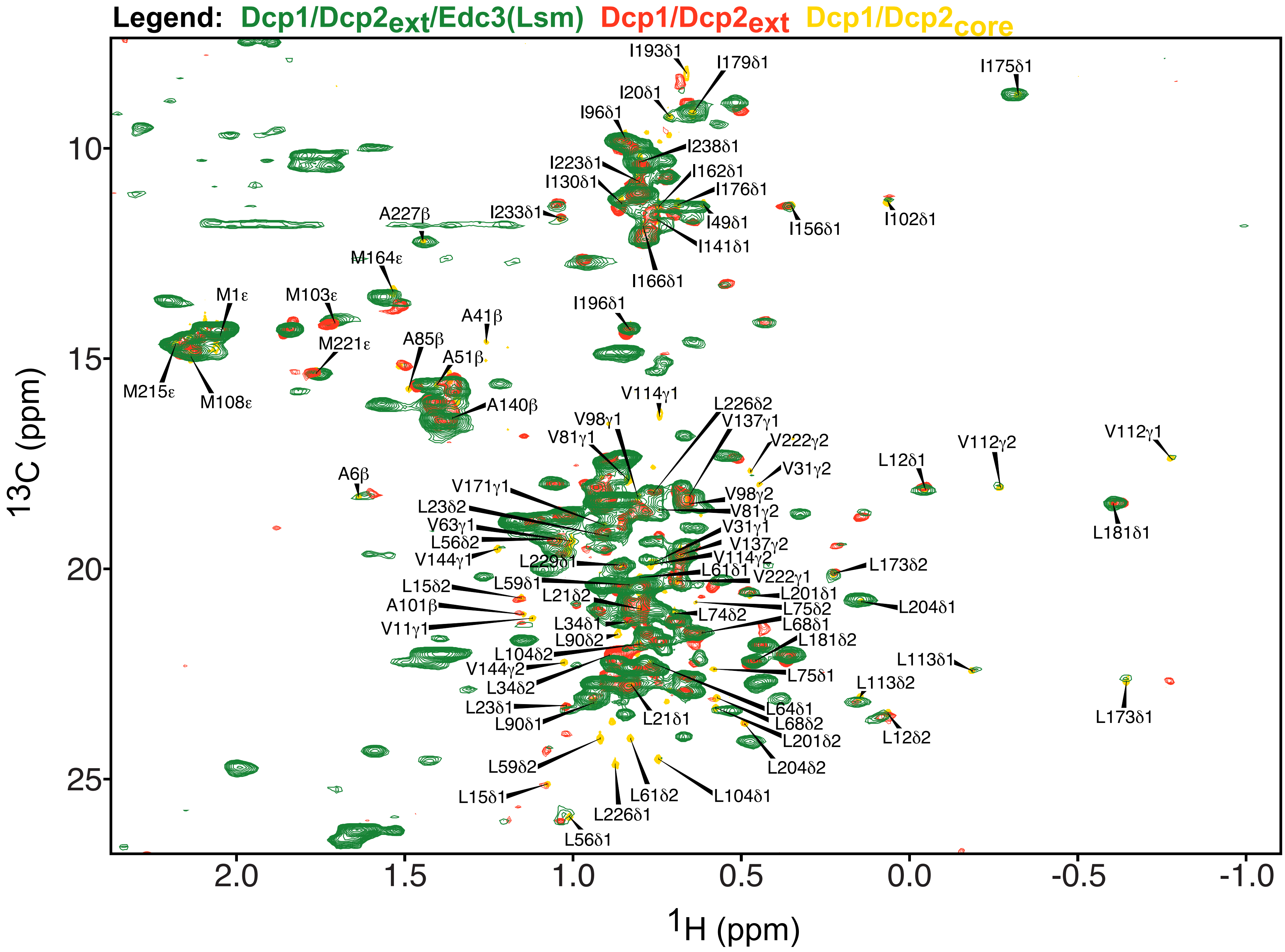

Legend: Dcp1/Dcp2ext/Edc3(Lsm) Dcp1/Dcp2ext Dcp1/Dcp2core
I193δ1
I179δ1
I175δ1
I20δ1
I96δ1
I238δ1
I223δ1
I162δ1
I130δ1
I176δ1
I49δ1
I156δ1
I102δ1
I141δ1
I233δ1
A227β
I166δ1
M164ε
M1ε
M103ε
I196δ1
A41β
M221ε
A85β
A51β
M215ε
M108ε
A140β
V114γ1
L226δ2
V137γ1
V98γ1
V81γ1
V222γ2
V31γ2
L12δ1
V112γ2
V112γ1
A6β
V171γ1
V98γ2
L23δ2
V81γ2
L181δ1
V63γ1
V31γ1
L56δ2
V137γ2
V144γ1
L229δ1
V114γ2
L61δ1
L173δ2
V222γ1
L15δ2
L59δ1
L21δ2
L201δ1
A101β
L204δ1
L75δ2
L74δ2
V11γ1
L34δ1
L68δ1
L90δ2
L181δ2
L104δ2
L113δ1
V144γ2
L75δ1
L34δ2
L113δ2
L23δ1
L64δ1
L21δ1
L173δ1
L90δ1
L68δ2
L201δ2
L12δ2
L59δ2
L204δ2
L61δ2
L104δ1
L15δ1
L226δ1
L56δ1
13C (ppm)
10
15
20
25
2.0
1.5
1.0
0.5
0.0
-0.5
-1.0
1H (ppm)

**Figure S8**

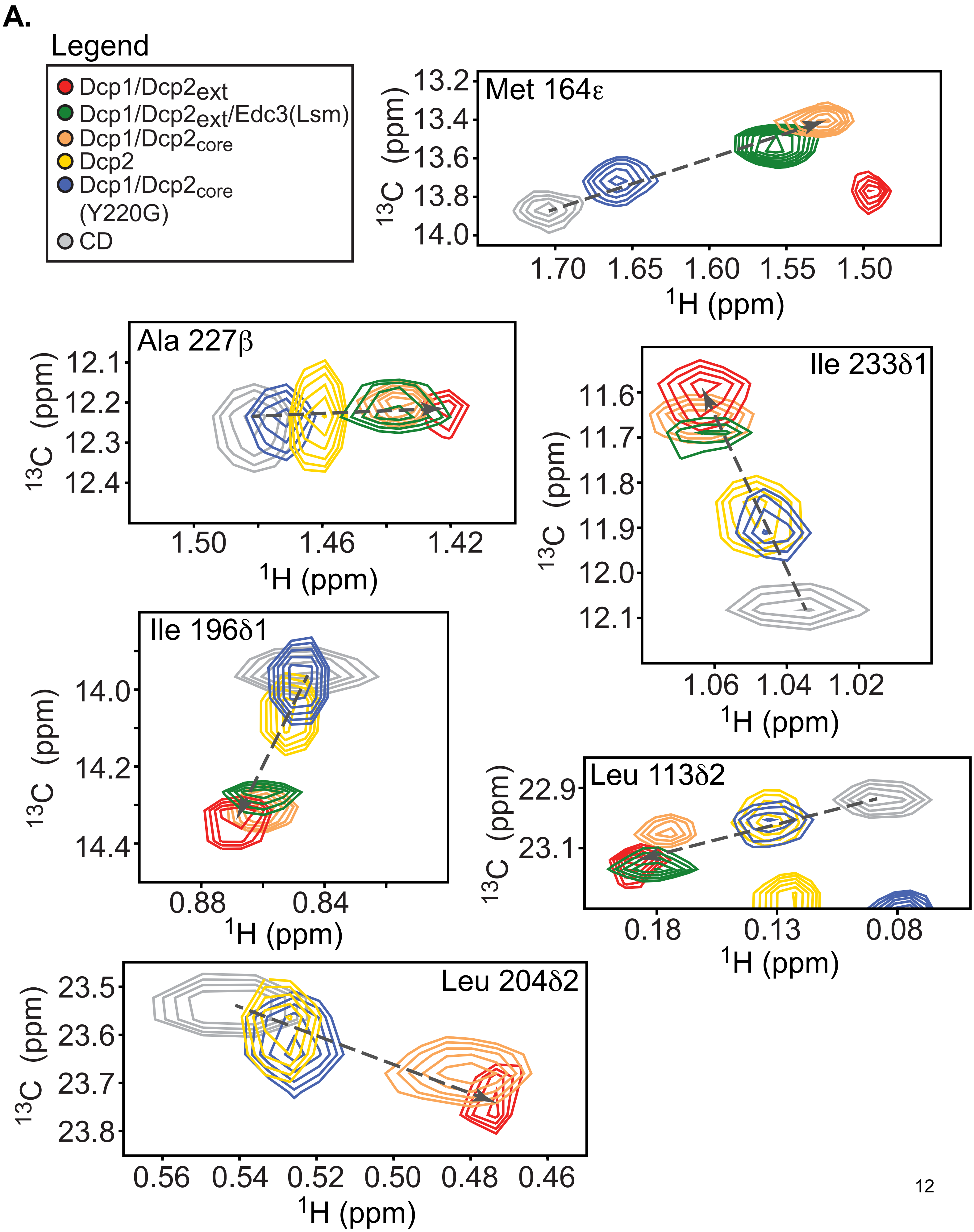
A.
Legend
Dcp1/Dcp2ext
Dcp1/Dcp2ext/Edc3(Lsm)
Dcp1/Dcp2core
Dcp2
Dcp1/Dcp2core (Y220G)
CD
Met 164ε
13.2
13.4
13.6
13.8
14.0
1.70
1.65
1.60
1.55
1.50
13C (ppm)
1H (ppm)
Ala 227β
12.1
12.2
12.3
12.4
1.50
1.46
1.42
Ile 233δ1
11.6
11.7
11.8
11.9
12.0
12.1
1.06
1.04
1.02
Ile 196δ1
14.0
14.2
14.4
0.88
0.84
Leu 113δ2
22.9
23.1
0.18
0.13
0.08
Leu 204δ2
23.5
23.6
23.7
23.8
0.56
0.54
0.52
0.50
0.48
0.46

**Figure S9**

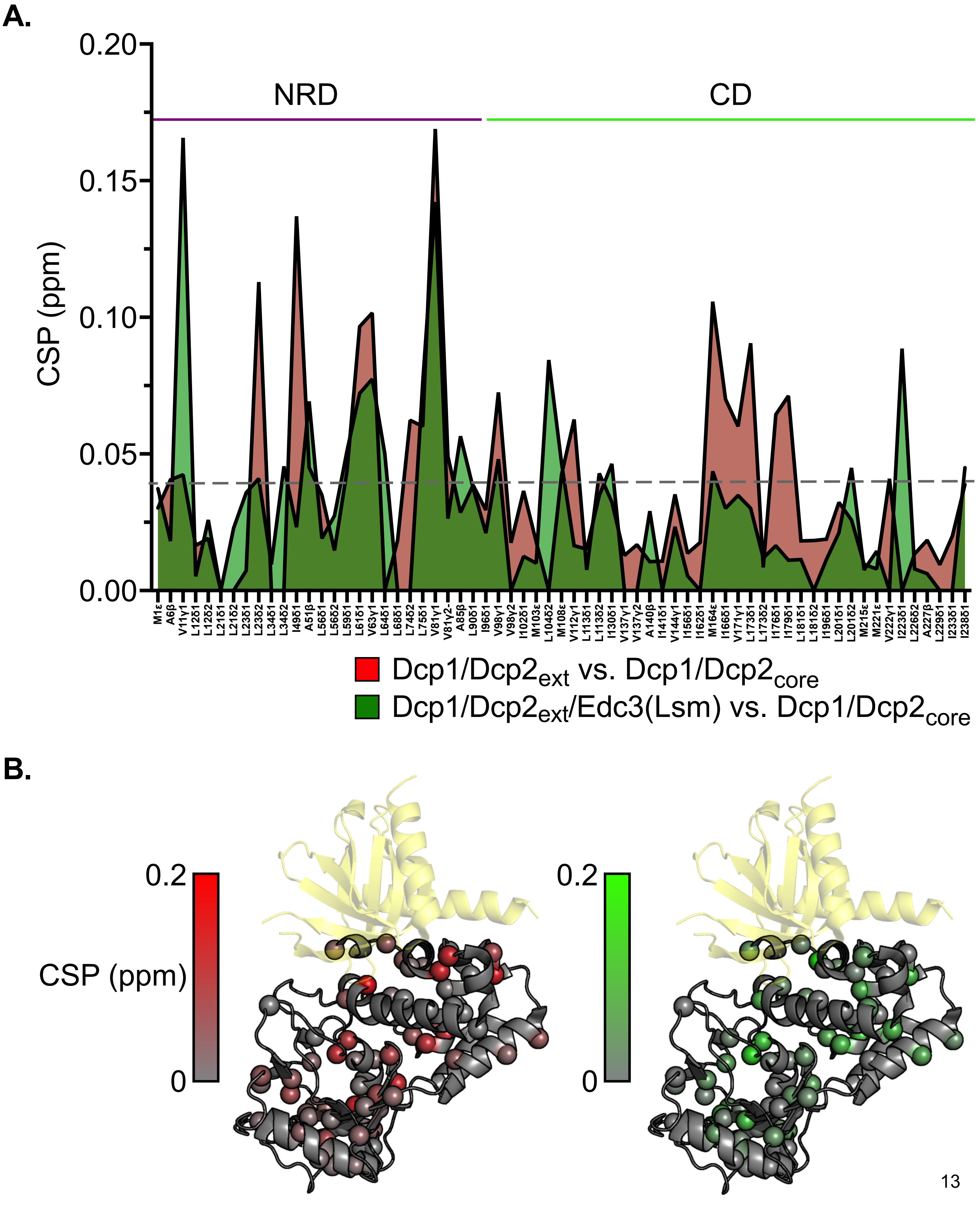
A.
0.20
0.15
0.10
0.05
0.00
CSP (ppm)
NRD
CD
Dcp1/Dcp2$_{ext}$ vs. Dcp1/Dcp2$_{core}$
Dcp1/Dcp2$_{ext}$/Edc3(Lsm) vs. Dcp1/Dcp2$_{core}$
B.
0.2
CSP (ppm)
0
0.2
0

#### Figure S10

A.

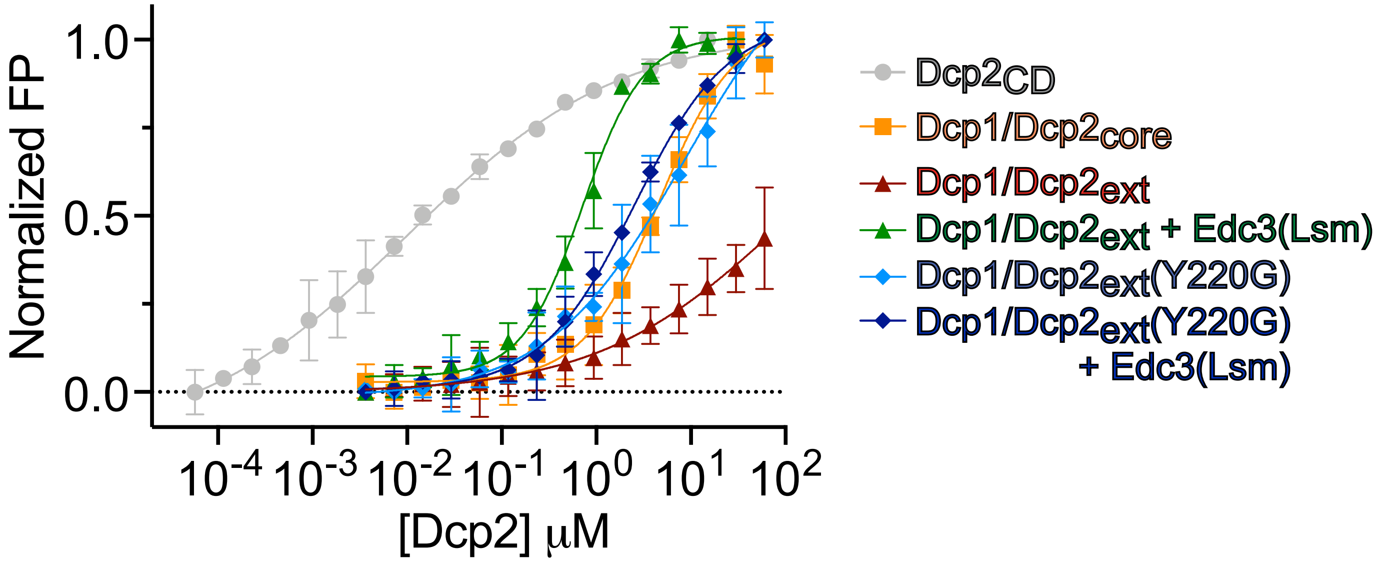


B.

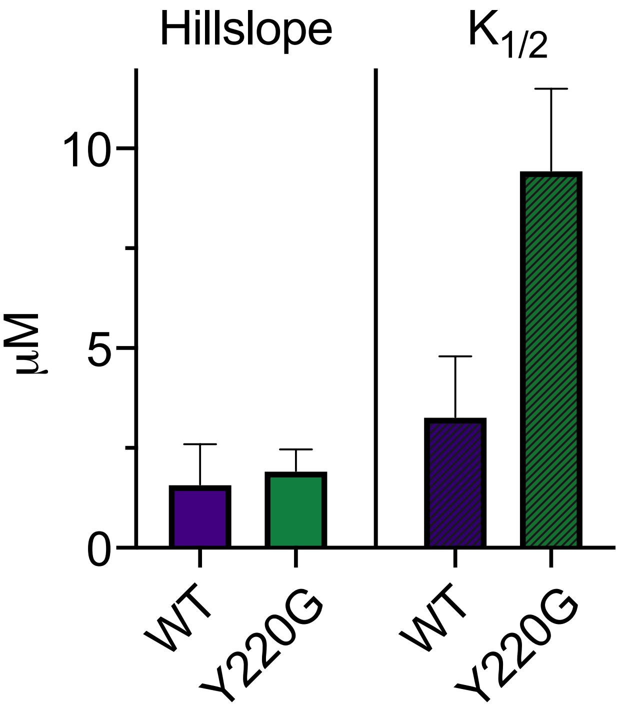

**Supplementary Table 1**: Protein constructs used in this study

| Protein Construct | Amino Acid Boundaries | Solubility/Purification Tags | Vector Backbone | Organism |
|---|---|---|---|---|
| Dcp1/Dcp2$_{ext}$ | Dcp1: 1-127<br>Dcp2: 1-504 | Dcp1: N-terminal His$_6$-MBP-TEV<br>Dcp2: C-terminal StrepII | pRSF | *S. pombe* |
| Dcp1/Dcp2$_{ext}$(Y220G) | Dcp1: 1-127<br>Dcp2: 1-504 | Dcp1: N-terminal His$_6$-MBP-TEV<br>Dcp2: C-terminal StrepII | pRSF | *S. pombe* |
| Dcp1/Dcp2$_{core}$ | Dcp1: 1-127<br>Dcp2: 1-243 | Dcp1: N-terminal His$_6$-MBP-TEV | pRSF | *S. pombe* |
| Dcp1/Dcp2$_{core}$(Y220G) | Dcp1: 1-127<br>Dcp2: 1-243 | Dcp1: N-terminal His$_6$-MBP-TEV | pRSF | *S. pombe* |
| Dcp1/Dcp2$_{HLM1/2}$ | Dcp1: 1-127<br>Dcp2: 1-318 | Dcp1: N-terminal His$_6$-MBP-TEV<br>Dcp2: C-terminal StrepII | pRSF | *S. pombe* |
| Dcp1/Dcp2$_{HLM1}$ | Dcp1: 1-127<br>Dcp2: 1-266 | Dcp1: N-terminal His$_6$-MBP-TEV | pRSF | |
| Dcp1/Dcp2(eSrtA) | Dcp1: 1-27<br>Dcp2: 1-266 + LPETGGH | Dcp1: N-terminal His$_6$-MBP-TEV | pRSF | *S. pombe* |
| Dcp1 | Dcp1: 1-127 | Dcp1: N-terminal His$_6$-MBP-TEV | pRSF | *S. pombe* |
| Dcp2$_{ext}$ | Dcp2: 1-504 | Dcp2: N-terminal His$_6$-MBP-TEV; C-terminal StrepII | pRSF | *S. pombe* |
| Dcp2$_{core}$ | Dcp2: 1-243 | Dcp2: N-terminal His$_6$-MBP-TEV | pRSF | *S. pombe* |
| Dcp2$_{CD}$ (<u>C</u>atalytic <u>D</u>omain) | Dcp2: 96-243 | Dcp2: N-terminal His$_6$-MBP-TEV | pRSF | *S. pombe* |
| Dcp2 C-terminus | Dcp2: 244-504 | Dcp2: N-terminal His$_6$-MBP-TEV; C-terminal StrepII | pRSF | *S. pombe* |
| Dcp2 C-terminus eSrtA | Dcp2: $G_3$ + 274-504 | Dcp2: N-terminal His$_6$-MBP-TEV; C-terminal StrepII | pRSF | *S. pombe* |
| Edc3 | Edc3: 1-454 | N-terminal His$_6$-TEV | pET30b | *S. pombe* |
| Edc3 Lsm domain | Edc3: 1-94 | N-terminal His$_6$-TEV | pET30b | *S. pombe* |

**Supplementary Table 2**: Relative activity of Dcp1/$Dcp2_{ext}$ in liquid droplets with or without Edc3

| Protein | Bulk Relative $k_{obs}$ ($min^{-1}$) | Supernatant Relative $k_{obs}$ ($min^{-1}$) |
|---|---|---|
| Dcp1/$Dcp2_{ext}$ | 0.71 ± 0.11 | 1 |
| Dcp1/$Dcp2_{ext}$/Edc3 | 3.19 ± 0.04 | 1 |

**Supplementary Table 3**: Relative activity of Dcp1/$Dcp2_{ext}$ in Bulk and Supernatant with variable Edc3 concentrations

| [Edc3] (μM) | Bulk Relative $k_{obs}$ ($min^{-1}$) | Supernatant Relative $k_{obs}$ ($min^{-1}$) |
|---|---|---|
| 80 | 3.03 ± 0.04 | 1.05 ± 0.03 |
| 20 | 3.08 ± 0.06 | 1.26 ± 0.1 |
| 5 | 2.6 ± 0.05 | 1.05 ± 0.08 |
| 1.25 | 1.31 ± 0.06 | 1.08 ± 0.03 |
| 0.3125 | 1.18 ± 0.02 | 1.18 ± 0.06 |
| 0.078125 | 1.09 ± 0.06 | 1.06 ± 0.04 |
| 0 | 1 | 1 |

**Supplementary Table 4**: Relative activity of Dcp1/$Dcp2_{ext}$ upon addition of varying Edc3 Lsm domain concentrations

| [Edc3 Lsm] (μM) | Relative $k_{obs}$ ($min^{-1}$) |
|---|---|
| 80 | 2.50 ± 0.19 |
| 20 | 2.14 ± 0.12 |
| 5 | 1.83 ± 0.11 |
| 1.25 | 1.42 ± 0.08 |
| 0.3125 | 1.16 ± 0.05 |
| 0.078125 | 1.01 ± 0.05 |
| 0 | 1 |

**Supplementary Table 5**: Relative activity of Dcp1/$Dcp2_{ext}$(Y220G) and Dcp1/$Dcp2_{ext}$ WT in Bulk and Supernatant at variable Edc3 concentrations

| [Edc3] (μM) | Y220G Bulk Relative $k_{obs}$ ($min^{-1}$) | Y220G Supernatant Relative $k_{obs}$ ($min^{-1}$) | WT Bulk Relative $k_{obs}$ ($min^{-1}$) | WT Supernatant Relative $k_{obs}$ ($min^{-1}$) |
|---|---|---|---|---|
| 80 | 10.03 ± 0.28 | 8.01 ± 0.16 | 3.11 ± 0.03 | 1.17 ± 0.04 |
| 20 | 9.41 ± 0.04 | 6.47 ± 0.32 | 2.82 ± 0.09 | 1.09 ± 0.06 |
| 5 | 7.39 ± 0.01 | 6.15 ± 0.42 | 2.45 ± 0.05 | 0.87 ± 0.07 |
| 1.25 | 6.59 ± 0.37 | 6.96 ± 0.02 | 1.32 ± 0.01 | 0.99 ± 0.03 |
| 0.3125 | 6.85 ± 0.55 | 7.36 ± 0.13 | 1.43 ± 0.09 | 1.01 ± 0.06 |
| 0.078125 | 6.39 ± 0.40 | 6.51 ± 0.58 | 0.88 ± 0.07 | 0.91 ± 0.05 |
| 0 | 6.42 ± 0.34 | 6.46 ± 0.25 | 1 | 1 |

**Supplementary Table 6**: Equilibrium dissociation constants ($K_D$) for various Dcp2 constructs determined from fluorescence polarization

| **Protein** | **$K_D$ ($\mu M$)** |
|---|---|
| $Dcp2_{CD}$ | 0.0097 ± 0.0031 |
| Dcp1/$Dcp2_{core}$ | 4.57 ± 1.04 |
| Dcp1/$Dcp2_{ext}$ | 84.1 ± 62.5 |
| Dcp1/$Dcp2_{ext}$ + Edc3(Lsm) | 0.837 ± 0.1207 |
| Dcp1/$Dcp2_{ext}$(Y220G) | 8.58 ± 3.69 |
| Dcp1/$Dcp2_{ext}$(Y220G) + Edc3(Lsm) | 2.53 ± 0.24 |
